# Task Engagement Gates Interareal Communication Geometry in the Mouse Thalamocortical–Midbrain Visual Circuit

**DOI:** 10.64898/2026.09.15.751810

**Authors:** Loris Amalberti, Maximilian Hauer, Corbett Bennett, Shawn R. Olsen, David Dahmen, Stefano Recanatesi

**Author notes:** These authors share senior authorship (David Dahmen, Stefano Recanatesi).

## Abstract

Visual processing unfolds across hierarchically organized brain circuits. Existing theories largely explain changes in population geometry through local shifts in gain, firing-rate statistics, or recurrent dynamics, yet do not account for how interareal coordination interacts with local population geometry to constrain downstream population states. We used task engagement, compared to a passive condition, to probe this coordination in Neuropixels recordings spanning the mouse visual thalamocortical–midbrain circuit. Engagement reduced network activity, response participation, and dimensionality across the hierarchy. To account for this circuit-level organization, we developed a theoretical framework in which afferent population geometry interacts with local recurrent dynamics to constrain the accessible dynamics of downstream populations. Across the thalamocortical stages, population-wide afferent statistics predicted downstream activity and dimensionality. At the cortex– midbrain interface, engagement instead reorganized interareal communication geometry. Together, these results identify interareal input geometry as a key constraint on neural population dynamics and uncover a general principle by which behavioral engagement constrains neural state spaces across distributed visual circuits.

## Introduction

Adaptive behavior emerges from coordinated computations distributed across hierarchically organized brain circuits, including thalamus, cortex, and midbrain (1, 2). Across these regions, local population dynamics are shaped by the interplay between recurrent connectivity and afferent drive, enabling flexible transformation of sensory information under changing behavioral demands. Additionally, behavioral engagement is known to strongly modulate these dynamics (3– 9). Yet, despite extensive characterization of state-dependent modulation at the level of firing rates and pairwise correlations (10–12), it remains unclear how engagement reshapes the geometry of neural population activity across interconnected areas at the circuit level.

Existing theoretical frameworks have largely characterized behavioral modulation from a local perspective, emphasizing how changes in gain, excitability, or recurrent stability reshape activity patterns within individual areas (10, 13). However, such local accounts are incomplete in an interconnected hierarchy, where structured afferent activity from upstream areas can itself constrain downstream population geometry. A central question therefore remains unresolved: how do engagement-dependent changes in afferent population statistics interact with local recurrent dynamics to reorganize neural population geometry across successive stages of a circuit?

To address this question, we focused on the mouse visual thalamocortical–midbrain hierarchy (1) (Fig. 1a), and we developed a theoretical framework describing how population statistics are transformed across these successive circuit stages (Supplementary Information, Sections 1–4). Whereas previous approaches have often emphasized either the effects of external drive or recurrent interactions (14–24), our framework expands these theoretical accounts by explicitly coupling structured afferent population geometry to the recurrent dynamics of downstream populations (Fig. 1b). In particular, our theory predicts that afferent variance and dimensionality exert largely dissociable effects on downstream population statistics. Structured afferent input can therefore expand or compress downstream dimensionality depending on the local operating regime, providing a direct link between interareal input statistics and local population geometry.

**Fig. 1.**
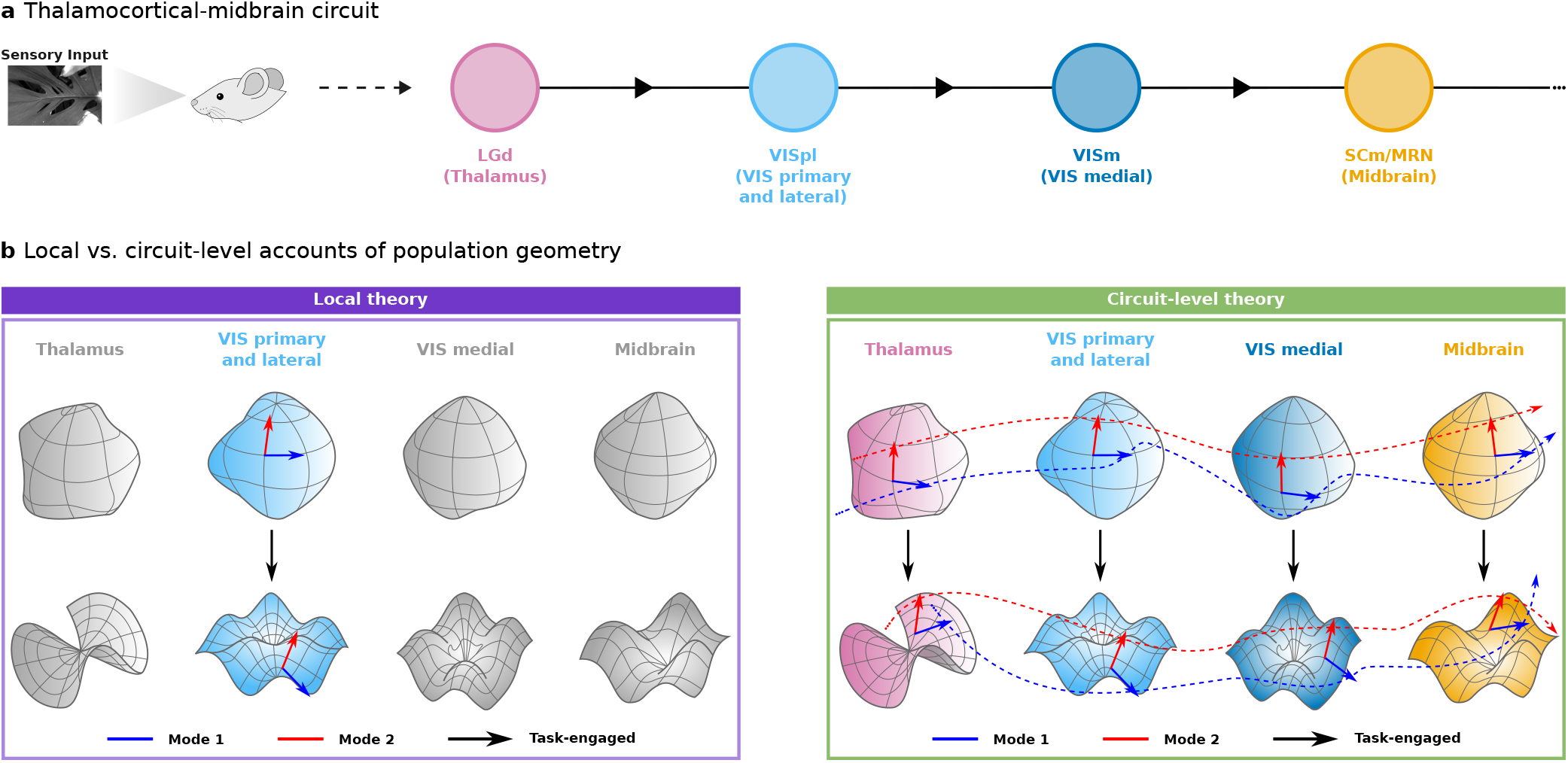
Local and circuit-level views of engagement-dependent population geometry. a) Schematic of the thalamocortical–midbrain hierarchy considered in this study (1). Visual input is relayed through the dorsal lateral geniculate nucleus (LGd) to primary and lateral visual cortex (VISpl: primary–VISp, rostrolateral–VISrl, lateral–VISl, anterolateral–VISal), medial visual cortex (VISm: posteromedial–VISpm, anteromedial–VISam), and downstream motor-related superior colliculus and midbrain reticular nucleus (SCm/MRN). b) Conceptual comparison of local and circuit-level accounts of population geometry. Population activity in each area is represented as a manifold in state space, with the colored modes providing a general coordinate system for population activity. In the local account, engagement-dependent reshaping and compression arise independently from local circuit dynamics within each area. In the circuit-level account, afferent population geometry (dashed trajectories) propagates through successive stages and interacts with local dynamics, thereby constraining the geometry of downstream population activity.

To test these predictions, we analyzed large-scale Neuropixels recordings spanning this hierarchy under two conditions: active engagement in a visual change detection task, in which mice reported image changes by licking for reward, and passive viewing of an identical stimulus sequence without the task requirement (1). Task engagement broadly reorganized population responses, reducing firing rates and response participation while compressing dimensionality relative to passive viewing (Fig. 1b). These changes were co-ordinated across interconnected areas rather than confined to a single stage of the hierarchy, consistent with a circuit-level reorganization of population responses. Across multiple circuit stages, source-area population statistics were associated with the matched statistics of downstream populations, in agreement with the theoretical prediction that structured afferent activity constrains downstream geometry. This mapping was nevertheless stage dependent, with the cortical– midbrain interface departing from the simple dissociation between variance and dimensionality predicted by the theory, indicating a more distinct organization of population geometry at this stage.

Together, these results identify engagement-dependent population geometry as a circuit-level phenomenon shaped by structured input statistics and interareal coordination, rather than a purely local property of individual regions. More broadly, our theoretical framework and empirical findings suggest that task engagement reorganizes the population subspaces through which information propagates across the thalamocortical–midbrain hierarchy, progressively constraining the state space accessible to downstream populations and providing a circuit-level mechanism for flexible sensory processing and behavior.

### Engagement compresses cortical population geometry

We analyzed neural activity from the Visual Behavior Neuropixels dataset recorded at the Allen Brain Observatory (25), which is based on a head-fixed visual change-detection task (Figures 2a to 2b; Fig. S1a). In this go/no-go paradigm, mice viewed a continuous stream of natural images presented for 250 ms and separated by 500 ms isoluminant gray screens. Within each trial, the same image was repeated a variable number of times before a different image appeared (the “change” stimulus). Mice were trained to report image changes by licking a water spout during a fixed post-change response window (150–750 ms after the change), with correct licks triggering delivery of a water reward.

**Fig. 2.**
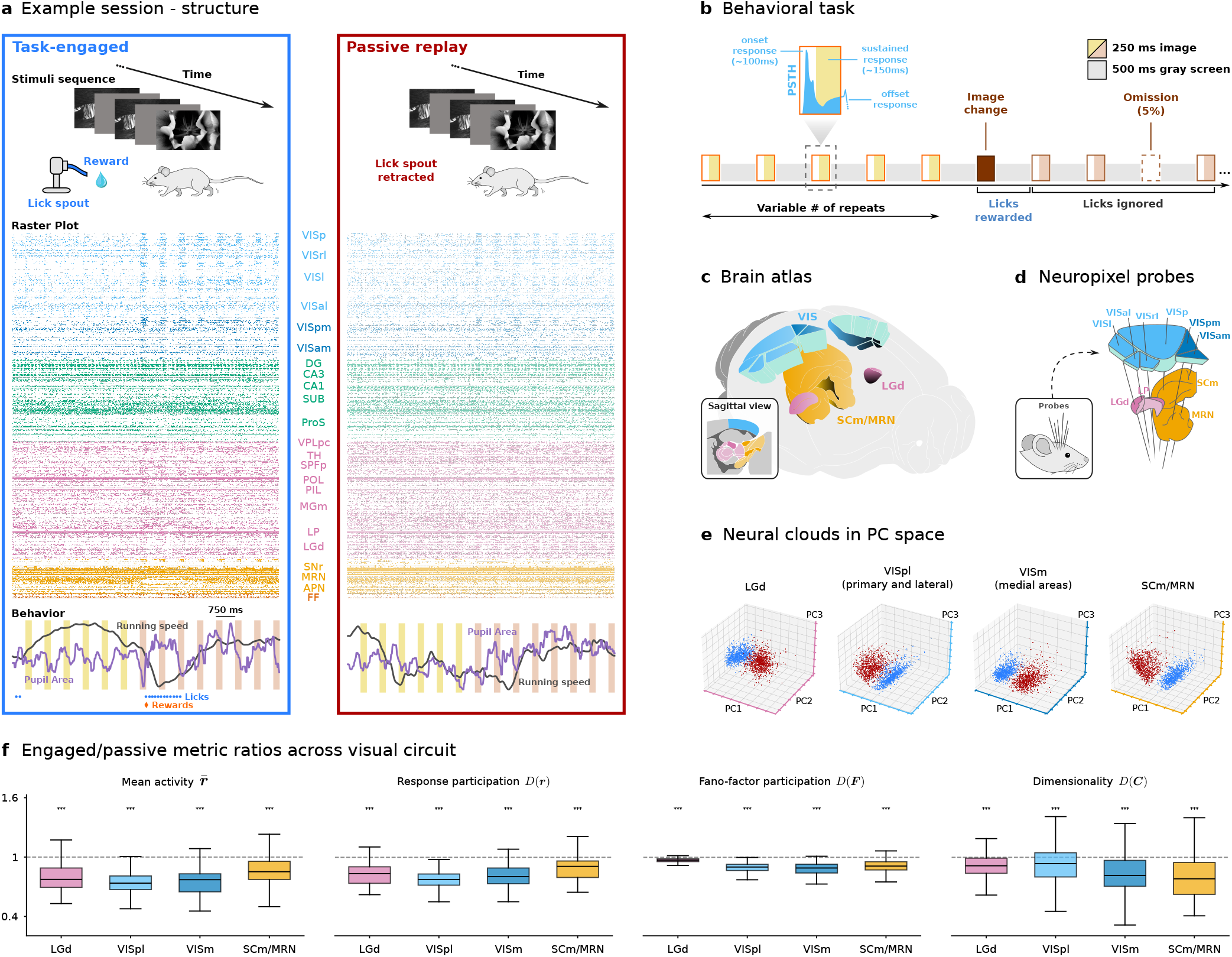
Task engagement reshapes population geometry across the visual thalamocortical–midbrain circuit. a) Example experimental session structure, illustrating active task engagement and passive replay blocks. Raster plots and behavioral variables are shown for an example session. Top, spike raster across recorded units. Bottom, running speed (dark gray), pupil area (purple), licks (blue), and reward delivery (orange). Shaded columns indicate repeated presentations of the same image; changes in column color denote transitions to a new image identity. b) Timeline of an example trial during the visual change-detection task, together with a schematic peri-stimulus time histogram (PSTH) of primary visual cortex (VISp) activity illustrating the early onset response (20–100 ms) and subsequent sustained response (100–250 ms; light yellow). Population analyses focused on the sustained response window of image repetitions to characterize the stable post-transient operating regime while excluding the enhanced activity associated with image changes. c) Anatomical localization of the recorded areas in a 3D brain atlas, highlighting visual cortical regions (VIS), the dorsal lateral geniculate nucleus (LGd), and the midbrain structures SCm and MRN. The inset shows the corresponding sagittal view. d) Schematic of Neuropixels probe placement across recorded brain regions. e) Population activity from a representative session and image projected onto the leading principal components for LGd, primary and lateral visual cortex (VISpl), medial visual cortex (VISm), and SCm/MRN during task engagement and passive viewing. f) Task-engaged/passive ratios of mean activity, response participation, Fano-factor participation, and dimensionality across LGd, VISpl, VISm, and SCm/MRN. Boxplots show session *×* image observations (8 images; *n* = 16 LGd, 48 VISpl, 47 VISm, and 31 SCm/MRN sessions); individual observations are shown in Fig. S4b. The dashed line marks a ratio of 1. Asterisks denote two-sided one-sample *t*-tests against 1: ^∗∗∗^*p <* 0.001.

During each recording session, neural activity was simultaneously recorded from multiple visual and subcortical brain areas (Figures 2c to 2d) during both active task performance and passive replay, in which the same stimulus sequence was presented without the lick spout or reward (Fig. 2a). Mice were trained on the stimulus set before data acquisition (Fig. S1b). We analyzed 48 sessions comprising large-scale population recordings (Fig. S1c). We further distinguished an early onset-response window (20–100 ms after image on-set (1)) from a subsequent sustained-response window (100– 250 ms), and focused our population analyses on the latter during image repetitions, because sustained responses more closely matched the activity regime described by our theory (Fig. 2b).

We first compared active engagement and passive viewing in primary visual cortex (VISp), the first task-involved cortical stage (1). Within the sustained-response regime, we described population activity by the vector of neuronal firing rates ***r*** = (*r*_1_, … , *r*_*N*_) (cf. Methods) and asked whether active and passive conditions occupied distinct regions of neural state space around identifiable operating points, defined as condition-specific averages of repeatable population activity patterns. In a low-dimensional projection obtained by Principal Component Analysis (PCA; cf. Methods), these operating points correspond to the centroids of the condition-specific response clouds. Active and passive response clouds were visibly separated across the recorded visual cortical areas (Figures S2a to S2b), indicating a widespread state-dependent shift in population activity.

In VISp, mean population activity E[***r***] was lower during active engagement than during passive viewing (Fig. S2c). Active engagement also significantly reduced response participation (Fig. S2c), as quantified by the Treves–Rolls metric (*D*(***r***) = E[***r***]^2^ */*E [***r***^2^]), where lower values correspond to sparser or less broadly distributed firing rates across neurons (26). To quantify the effective dimensionality of population activity, we measured the participation ratio (*D*(***C***) = E[***λ***]^2^ */*E[***λ***^2^]) of the covariance matrix ***C***, where *λ*_*i*_ are its *N* eigenvalues obtained via PCA. Task engagement reduced *D*(***C***) (Fig. S2c), indicating that trial-to-trial variability was concentrated into fewer population modes. Thus, engagement simultaneously reduced mean activity and response participation while compressing population dimensionality. These effects extended to lateral visual cortical areas (VISrl, VISl, VISal) and medial visual cortical areas (VISpm, VISam) (Fig. S2c) and were consistent across individual stimuli (Fig. S2d). We also asked whether similar changes occurred as engagement fluctuated during the active task, using licking patterns to distinguish engaged from unengaged periods (Figures S3a to S3b). Across visual cortical areas, mean activity and response participation during unengaged periods became more similar to those observed during passive viewing, but dimensionality did not differ significantly between engaged and unengaged periods (Fig. S3c). This suggests that the dimensionality difference between active task performance and passive viewing may reflect broader differences between these conditions, beyond fluctuations in engagement inferred from licking.

Together, these results show that task engagement reconfigures population dynamics and compresses population geometry across visual cortex. This raises a mechanistic question: can this compression be explained by local changes in cortical dynamics, such as altered recurrent gain or intrinsic variability, or does it reflect engagement-dependent changes in afferent drive across the visual circuit?

### From local models to circuit-level input geometry

The engagement-dependent changes described above recurred across individual visual cortical areas, suggesting a broader coordinated reorganization of local population dynamics that may extend across successive stages of the visual processing hierarchy. Distinguishing local from circuit-level mechanisms, however, requires first asking what local theories can, and cannot, explain. Existing theories of neural population geometry provide a natural first account: behavioral state can alter the local operating regime through changes in recurrent amplification, intrinsic variability, and neuronal gain (5, 19, 22, 24, 27–30). However, these accounts are intrinsically local and do not specify how structured afferent activity from upstream or interconnected areas is transformed into the covariance geometry of a downstream population (Fig. 1b). To make this local prediction explicit, we derived an expression relating dimensionality to other network-level observables (Supplementary Information, Section 5):

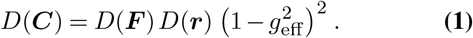

Here, *D*(***r***) denotes response participation, *D*(***F***) Fano-factor participation across neurons (Fig. S4a), and 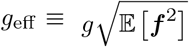 combines recurrent coupling strength *g* with the local neuronal gains *f*_*i*_ evaluated at the operating point (Supplementary Information, Section 1). This expression recovers the result of Tian et al. (30) in the limit of Poisson-like neurons and a quadratic threshold-power-law transfer function (Supplementary Information, Section 5).

To test whether this local account generalized across successive circuit stages, we extended the population-geometry analyses of Fig. S2 to siymultaneously recorded populations along the thalamocortical–midbrain circuit: LGd, VISpl, VISm, and SCm/MRN (1). Low-dimensional PCA projections revealed distinct active and passive population states at each stage (Fig. 2e). Across sessions, task engagement consistently reduced mean activity, response participation, and Fano-factor participation, as well as population dimensionality, in all four populations (Fig. 2f). The reductions in response participation and Fano-factor participation were consistent with the dimensionality reduction predicted by our local model. Active engagement also reduced analytically estimated neuronal gain moments across the hierarchy (Fig. S4c; Supplementary Information, Section 5). Together, these observations show that task engagement systematically shifts local operating regimes and compresses population geometry across successive stages of the circuit (Fig. S4d). These geometric effects were robust to temporal nonstationarity: dimensionality remained lower during engagement across successive within-condition time-slices (Figures S5a to S5b).

The recurrence of the same engagement-dependent changes across successive connected populations suggests that population geometry may be coordinated across the hierarchy. This could be captured by a circuit-level account in which engagement reshapes the statistics of activity propagated between areas, thereby constraining the operating regime of downstream populations. Such an analysis would provide a possible mechanistic link between the changes observed at successive stages and motivate a framework that explicitly couples afferent input geometry to local recurrent dynamics. We next introduce a theory formalizing this interaction and examine whether it can account for the population changes observed across the circuit.

### Input geometry constrains recurrent dynamics

We therefore formulate, building on previous accounts (19, 24), an input-explicit model that asks how structured afferent covariance is transformed by a recurrent target network. The model provides a tractable framework for isolating how input geometry, recurrent dynamics, and intrinsic noise jointly shape target population geometry (Supplementary Information, Sections 1–2). We model the target area as a recurrent neural network linearized around a stable operating point. The effective recurrent connectivity is evaluated at that operating point, thereby incorporating both anatomical coupling and neuron-specific gains. Source-area fluctuations enter the target dynamics through an effective afferent coupling matrix ***A***:

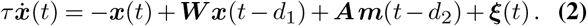

where *x*_*i*_(*t*) is the fluctuation of target neuron *i* around its operating point *r*_*i*_. The matrix *W*_*ij*_ is the effective recurrent connectivity after linearization, with neuronal gains absorbed into the recurrent couplings. *τ* is the neuronal time constant, the delays *d*_1_ and *d*_2_ denote recurrent and afferent transmission delays, respectively, and *ξ*_*i*_(*t*) captures additional unstructured intrinsic noise. The variables *m*_*α*_(*t*) (*α* ∈ {1, … , *M}*) are zero-mean source-area fluctuations expressed in the eigenbasis of the source covariance. Thus, source modes *m*_*α*_ are independent, each carrying variance *λ*_*α*_. Finally, the matrix entries *A*_*iα*_ are the effective afferent couplings from source covariance mode *α* to target neuron *i*, after gain rescaling and rotation into the source-covariance eigenbasis (Supplementary Information, Sections 1, 4). When the source covariance has low effective dimensionality, only a limited number of source modes dominate this term, imprinting the geometry of afferent drive onto the target covariance.

The output of the theory is a beyond-mean-field result of the higher-order activity moments needed to predict the target neural population statistics, including its dimensionality. The major advancement of this theory is that it allows tracking how structured source modes are mapped into the target population by the effective afferent coupling matrix, mixed by recurrent dynamics, and expressed in the second-order statistics of target activity. We discuss the full derivation in the Supplementary Information (Sections 3–4) and report here the central closed-form result:

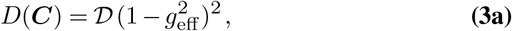

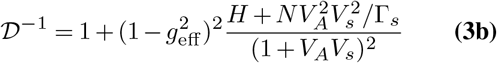

Here, *H* reflects heterogeneity of intrinsic fluctuations, *V*_*A*_ = Var(*A*_*iα*_) is the variance of the effective afferent coupling coefficients, *V*_*s*_ is the rescaled total source variance, and Γ_*s*_ is the source dimensionality (Supplementary Information, Section 4). Eq. (3a) can be regarded as a generalization of Eq. (1): the factor *D* reduces to the product *D*(***F***)*D*(***r***) in the absence of structured afferent input, i.e. for *V*_*s*_ = 0, and under assumptions of renewal spiking statistics (Supplementary Information, Section 5). Furthermore, for homogeneous single-neuron statistics *H* = 0 and *V*_*s*_ = 0, the factor simplifies to *D* = 1, which recovers the known result in (21, 24).

The central new implication of Eq. (3a) is that afferent drive constrains target geometry through source dimensionality, source variance, and the variance of the effective afferent couplings. Source dimensionality determines how many independent directions are injected into the target population: when source fluctuations are distributed across many modes, *i*.*e*., when Γ_*s*_ is high, the target receives more independent directions and its dimensionality increases. By contrast, the effect of increasing *V*_*s*_ depends on *V*_*A*_ and can either compress or expand target dimensionality (Fig. 3a). Thus, input dimensionality and variance exert distinct effects, with the influence of source variance modulated by the variance of the effective afferent couplings. Finally, the theory predicts a partial dissociation: source variance can constrain target dimensionality, whereas source dimensionality should not directly determine target variance. Across the hierarchy, empirical Fano-factor distributions were narrowly concentrated near the Poisson expectation of 1 (Fig. S4a), making spike-count variance proportional to mean activity, quantified here by the population-averaged firing rate. For this reason, we focus below on mean activity. The theory therefore yields two matched source–target predictions: higher source activity should predict higher target activity, whereas higher source dimensionality should predict higher target dimensionality (Fig. 3b).

**Fig. 3.**
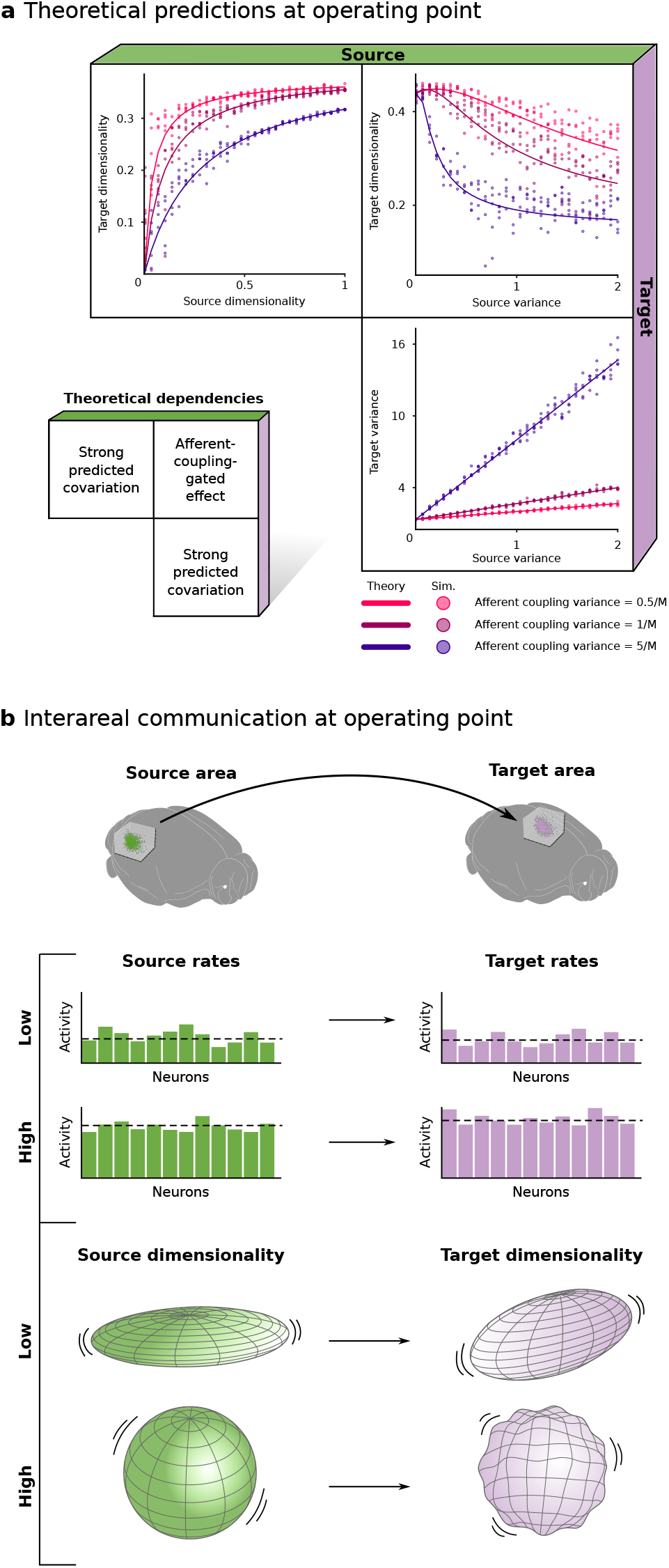
Input geometry constrains target-population activity and dimensionality. a) Theoretical predictions for how source dimensionality, source variance, and afferent coupling variance (*V*_*A*_) shape target statistics. Target dimensionality increases with source dimensionality, while source variance can either compress or expand it; target variance increases with source variance. Curves show theoretical expectations for *V*_*A*_ = 0.5*/M* , 1*/M* , and 5*/M* , with *M* = 1000 source modes; points show simulations. b) Schematic of interareal communication at the operating point. Higher source activity predicts higher target activity, and higher source dimensionality predicts higher target dimensionality.

### Source–target relationships are stage dependent

We next tested whether the input-dependent constraints predicted by the model were expressed across the recorded visual circuit, asking whether source-area population statistics were associated with the operating regime of downstream targets.

We tested these predictions along three successive source–target channels spanning the recorded hierarchy: LGd →VISpl, VISpl→VISm, and VISm→SCm/MRN (Fig. 2c). For each channel, we fit regression models predicting target mean activity or target dimensionality from source-area activity and source-area dimensionality. Because behavioral state can modulate activity in visual cortex and mid-brain, running speed and pupil area were included as additional covariates (31). This allowed us to assess source–target associations after adjustment for these measured locomotion- and arousal-related variables.

The source–target relationships were stage dependent (Figures 4a to 4b). Source mean activity was associated with target mean activity along all three channels. Source dimensionality, by contrast, was associated with target dimensionality at the LGd→VISpl and VISpl→VISm stages, but not significantly at VISm→SCm/MRN in either the active or passive block, where its contribution was subleading relative to other predictors (Figures 4b to 4c).

**Fig. 4.**
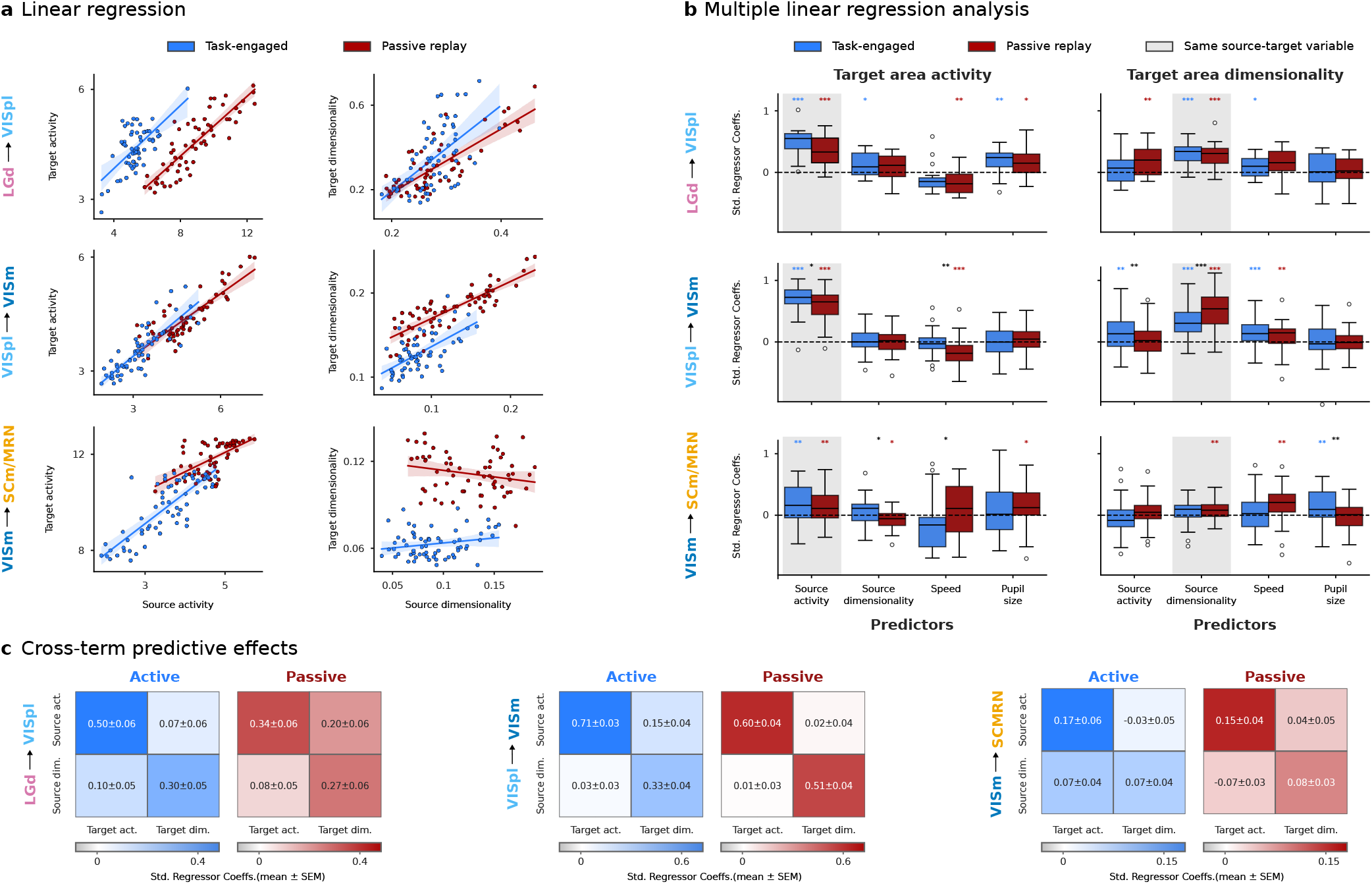
Source activity and dimensionality predict matched downstream population statistics. a) Example within-session regressions along the LGd→VISpl, VISpl→VISm, and VISm→SCm/MRN source–target orderings, shown separately for active and passive blocks. Each point is one image *×* 200-bin subsample; lines are ordinary least-squares fits with 95% confidence intervals. b) Standardized coefficients from models predicting target mean activity or dimensionality from source mean activity, source dimensionality, running speed, and pupil area. Boxplots show session-level coefficients (*n* = 16 LGd→VISpl, 47 VISpl→VISm, and 31 VISm→SCm/MRN sessions). Colored asterisks denote two-sided one-sample *t*-tests against zero; black asterisks denote two-sided paired *t*-tests between active and passive coefficients. ^∗^*p <* 0.05, ^∗∗^*p <* 0.01, ^∗∗∗^*p <* 0.001; *p* values are unadjusted. c) Schematic summary of source-activity and source-dimensionality coefficients from b; values report mean *±* SEM across sessions.

Because active and passive blocks also differed in locomotion and arousal-related variables (Fig. S3), we asked whether running speed or pupil area could account for the engagement-dependent dimensionality difference. As a complementary control, presentations were grouped by running-speed terciles (slow, moderate, fast) or pupil-area terciles (small, moderate, large), defined separately within each active and passive block (Fig. S6a). Dimensionality varied across these behavioral bins, with the increase at higher running speeds consistent with recent reports of higher cortical dimensionality during locomotion (32). We therefore fit an additional multiple-regression model including the active/passive condition together with condition-specific running-speed and pupil-area terms (Fig. S6b). The active/passive condition remained a significant predictor of dimensionality, whereas the running-speed and pupil-area terms were neither significant nor leading predictors. Thus, the active–passive dimensionality difference was not accounted for by these measured locomotion- and pupil-related variables.

Taken together, the regressions revealed a stage-dependent organization of source–target population statistics. Source and target mean activity covaried across all three channels, whereas dimensionality covaried across the thalamocortical and corticocortical stages but not detectably at the VISm→SCm/MRN transition. This departure does not necessarily imply an absence of structured cortex-to-midbrain coupling; instead, source dimensionality alone may be insufficient to describe the relevant afferent structure at this stage. Coupling may depend on which patterns of VISm activity covary with SCm/MRN activity, rather than simply on the overall dimensionality of the source population as considered in our model. We therefore identified these patterns of shared variation and asked whether they changed with engagement.

### Task engagement reorganizes inter-area covariance geometry

We focused on the geometry of VISm–SCm/MRN covariability directly and computed mean-centered local covariance matrices within each area and cross-covariance matrices between areas, separately for active engagement and passive viewing (Fig. 5a). We decomposed the VISm– SCm/MRN cross-covariance matrix using singular value decomposition (SVD), which identifies paired source and target activity modes that account for shared covariance across areas (Fig. 5b). Motivated by previous work (33, 34), we refer to the paired subspaces spanned by these SVD modes as the inter-area communication channel, and to the corresponding source- and target-side subspaces as communication sub-spaces. The leading singular values were larger during active engagement, and the active/passive ratios confirmed that the strongest coupled modes were enhanced in the active condition (Fig. 5c). Thus, engagement strengthened VISm– SCm/MRN covariability along its dominant shared modes.

**Fig. 5.**
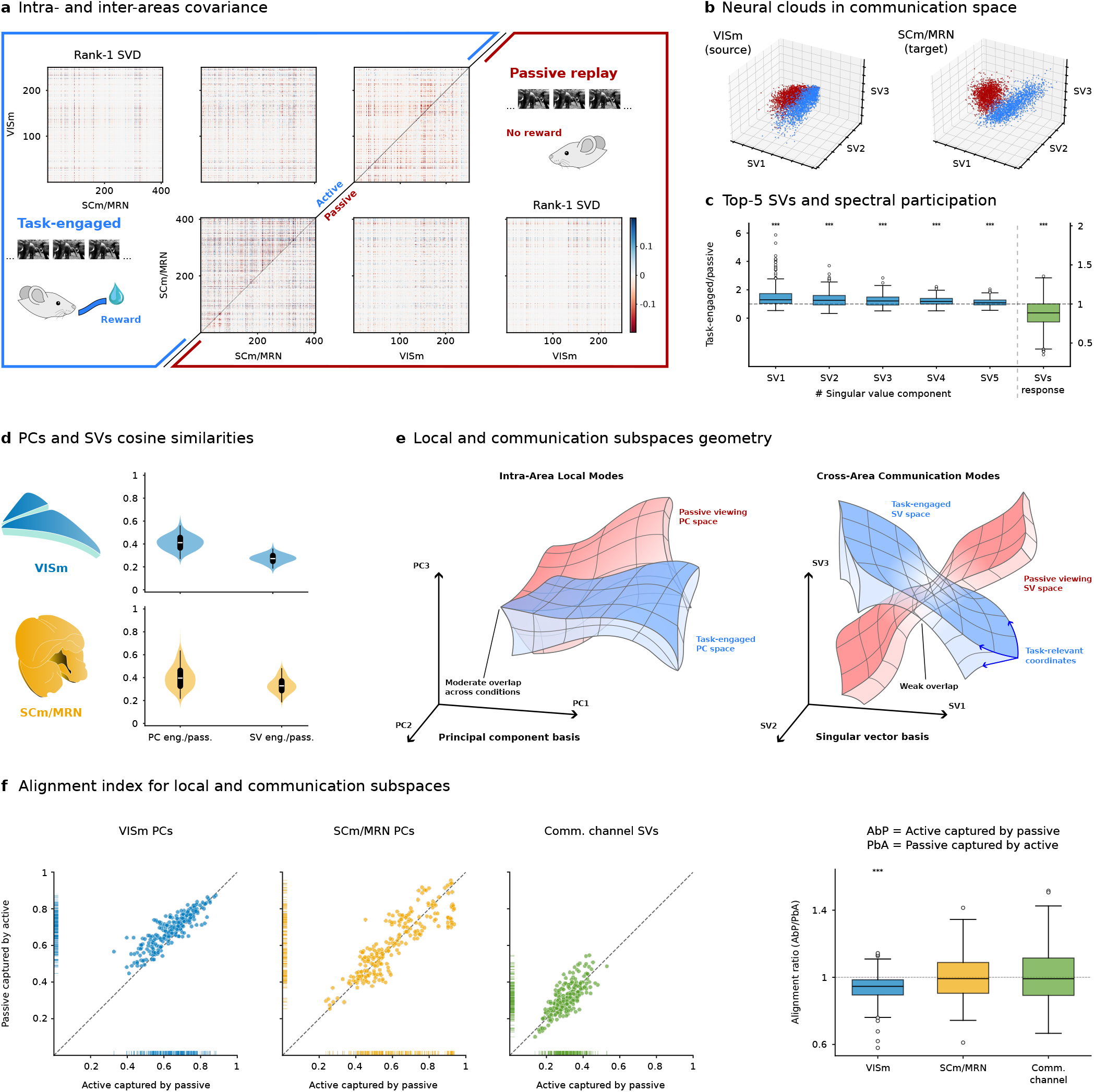
Active and passive blocks differ in VISm–SCm/MRN cross-covariance geometry. a) Mean-centered local covariance matrices for VISm and SCm/MRN and the corresponding VISm–SCm/MRN cross-covariance matrices from a representative session and image, shown for active and passive blocks. Rank-1 SVD reconstructions illustrate the leading cross-area covariance pattern. b) Population activity from the same representative session and image projected onto the leading source- and target-side singular-vector coordinates of the VISm–SCm/MRN cross-covariance matrix. c) Left, task-engaged/passive ratios of the five leading singular values of the VISm–SCm/MRN cross-covariance matrix. Right, the ratio of dimensionality computed from all singular values; values below 1 indicate that cross-area covariance is concentrated into fewer effective modes during engagement. Boxplots show session *×* image observations (*n* = 31 sessions, 8 images). Asterisks denote two-sided one-sample *t*-tests against 1: ^∗∗∗^*p <* 0.001. d) Mean absolute cosine similarity between task-engaged and passive modes for the leading *k* = 5 local principal components and cross-area singular vectors. Modes were paired across conditions by maximizing total absolute cosine similarity with the Hungarian algorithm. Violin plots show session *×* image observations (*n* = 31 sessions, 8 images). e) Conceptual comparison of local and cross-area population geometry. Dominant local PC subspaces remain partially overlapping across active and passive blocks, whereas the paired SVD-defined cross-area subspaces show weaker overlap across conditions. f) Directional cross-condition capture for the leading *k* = 5 local PC subspaces in VISm and SCm/MRN and the paired VISm–SCm/MRN communication subspaces. Scatter plots compare active covariance captured by the passive subspace (AbP) with passive covariance captured by the active subspace (PbA). Right, AbP/PbA; a ratio of 1 indicates symmetric capture. Boxplots show session *×* image observations as in c. Asterisks denote two-sided one-sample *t*-tests against 1: ^∗^*p <* 0.05, ^∗∗^*p <* 0.01, ^∗∗∗^*p <* 0.001.

We next asked whether engagement also changed the dimensionality of this communication structure. The dimensionality of the singular-value spectrum was significantly lower during active engagement than during passive viewing (Fig. 5c), indicating that shared covariance was concentrated into fewer effective modes. Thus, active engagement strengthened VISm–SCm/MRN cross-area covariance while concentrating it into fewer dominant modes, consistent with a lower-dimensional inter-area communication geometry.

We next tested whether the engagement-dependent covariance structure reflected a reorganization of local population geometry or a more selective change in inter-area communication modes. For each session, we compared active and passive conditions by matching the top 5 local PCs in VISm and SCm/MRN, as well as the top 5 paired SVD communication modes. Matches were obtained with a Hungarian algorithm that maximized the total absolute cosine similarity across modes, allowing for changes in mode ordering between conditions (cf. Methods).

This analysis revealed comparatively stronger active engagement–passive viewing alignment for local PCs than for cross-area SVD modes (Figures 5d to 5e). Local covariance axes were more strongly preserved across conditions, whereas the coupled VISm–SCm/MRN modes showed comparatively weaker alignment and greater state specificity. These results suggest that engagement more strongly reorganizes the cross-area communication modes than the dominant local covariance axes. An analogous VISpl–SCm/MRN analysis yielded the same qualitative pattern, with local PCs showing stronger active engagement–passive viewing alignment than cross-area SVD modes (Fig. S7a).

To further quantify state-dependent differences in communication geometry, we computed a cross-condition alignment index (35) (cf. Methods). This index measures, directionally, how much the covariance structure observed in one condition is captured by the subspace defined in the other condition. For the top-5 local PC subspaces, active–passive overlap was moderate to high in both VISm and SCm/MRN (Fig. 5f), consistent with the mode-wise cosine-similarity analysis. By contrast, the cross-area communication subspaces showed substantially weaker overlap across conditions (Fig. 5f), indicating greater state specificity of the communication modes.

This PC alignment index also revealed a directional asymmetry in VISm, but not SCm/MRN (Fig. 5f, right; Fig. S7b yields same qualitative pattern for VISpl–SCm/MRN analysis). Therefore, passive covariance was captured more strongly by the active PC subspace than active covariance was captured by the passive PC subspace (Fig. S7c). This asymmetry is consistent with task engagement preserving much of the passive covariance geometry while additionally recruiting high-variance directions that are weaker or absent during passive viewing.

Together, these analyses show that task engagement differentially affects local and inter-area covariance geometry. Dominant local PC subspaces remained more strongly overlapping across active and passive states, with a directional asymmetry in VISm, whereas leading cross-area communication subspaces showed comparatively weaker active–passive alignment. Engagement also concentrated shared covariance into fewer dominant communication modes. The cortex-to-midbrain transition is therefore better described as a state-dependent transformation of communication geometry than as simple propagation of dimensionality. This raised the next question: what task-related activity can be decoded from the communication dimensions defined during engagement?

### Communication subspaces capture lick-related activity differently in cortex and midbrain

The preceding analyses showed that engagement reorganizes the population dimensions along which VISm and SCm/MRN activity covary. We next asked whether these communication dimensions capture activity related to the animal’s behavior. Specifically, is lick-related activity preferentially represented in the communication subspace, or is it broadly accessible across other dimensions of each population? Addressing this question would clarify the behavioral relevance of the engagement-dependent changes in cross-area covariance.

We decoded lick versus no-lick labels from active-block population activity in individual 50-ms bins during non-change image presentations (Fig. 6a). Restricting the analysis to non-change presentations preserved the repeated-image response regime used throughout the population analyses. Lick-labeled samples therefore corresponded to non-rewarded licks outside the rewarded post-change response window, rather than correct change-detection reports or hits (Fig. S1a; see Methods). The analysis thus assesses activity associated with licking during task performance, rather than successful stimulus detection.

**Fig. 6.**
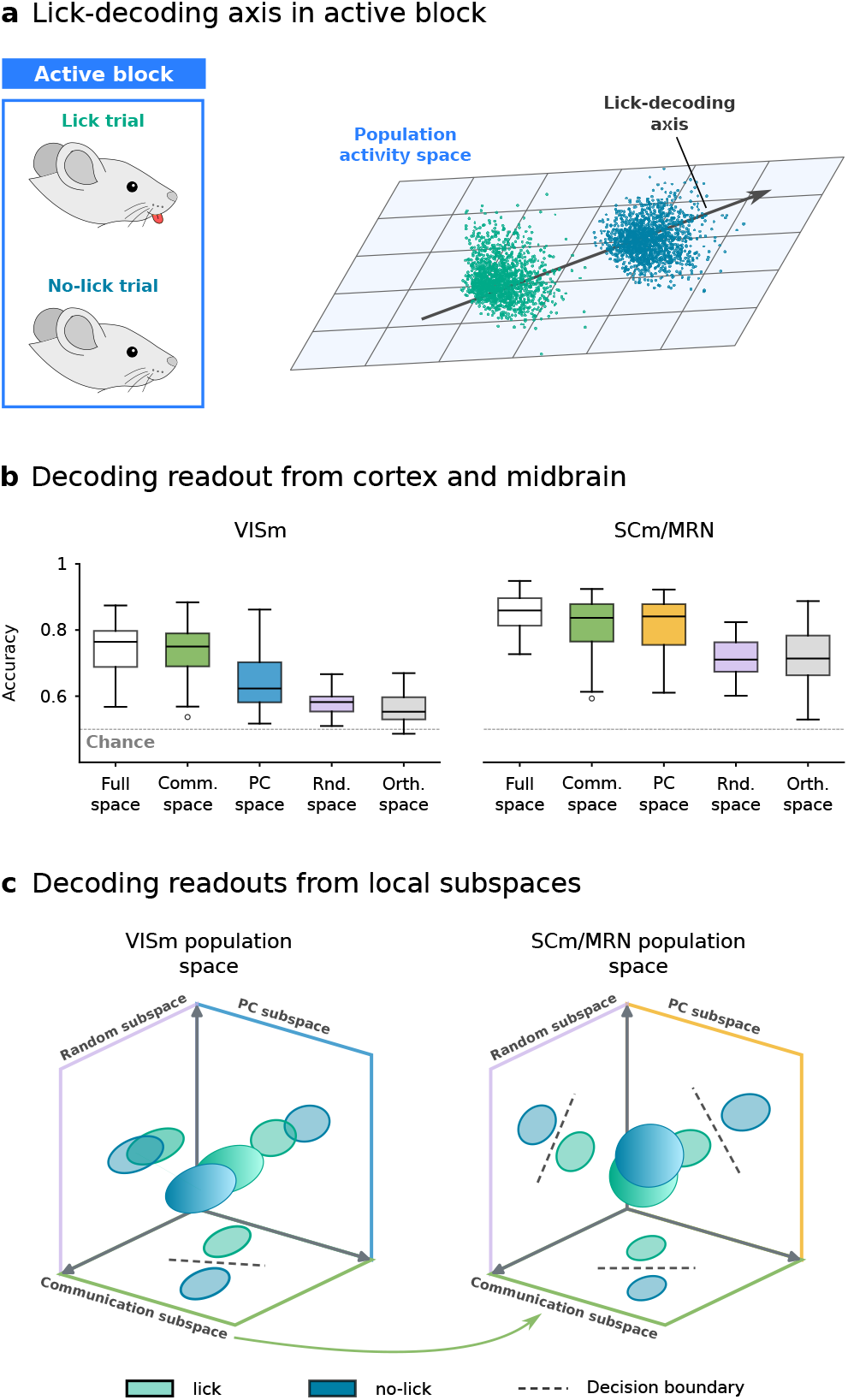
Lick-related activity is preferentially decoded from cortical communication dimensions and more broadly decoded in midbrain. a) Schematic of lick versus no-lick decoding from active-block population activity in individual 50-ms bins. Lick labels correspond to non-rewarded error licks during non-change image presentations, outside the rewarded post-change response window, rather than correct change-detection reports. b) Balanced accuracy in VISm and SCm/MRN (*n* = 31 sessions), comparing the full population space, the active communication subspace (*k* = 5), the active local-PC subspace (*k* = 5), matched random five-dimensional subspaces, and the communication-orthogonal complement. Performance was estimated using five-fold stratified cross-validation, with the decoder trained on four folds and evaluated on the held-out fifth fold. Boxplots show session-level means; the dashed line indicates chance (0.5). In VISm, communication dimensions nearly recover full-space decoding, whereas the orthogonal complement supports little decoding. In SCm/MRN, substantial decoding persists across other subspaces. c) Schematic interpretation of decoding across population sub-spaces in VISm (left) and SCm/MRN (right). Three-dimensional clouds represent full-population activity; colored planes represent communication, local-PC, and random subspaces. Greater separation of lick and no-lick activity across a linear decision boundary indicates better decoding. This separation is preferentially captured by communication dimensions in VISm and is apparent across multiple subspaces in SCm/MRN.

To determine how this activity was distributed within each population, we compared decoding from the full population space with decoding from three five-dimensional sub-spaces: the active communication subspace, which captures cross-area covariation; the active local-PC subspace, which captures the greatest within-area variance; and matched random subspaces. These comparisons test whether communication dimensions capture lick-related activity beyond what is expected from their dimensionality or from selecting high-variance local dimensions. We also decoded from the communication-orthogonal complement to determine how much lick information remained accessible outside the communication subspace.

In VISm, the five active communication dimensions supported nearly the same lick decoding as the full population: balanced accuracy was 0.73 *±*0.01, compared with 0.75 *±*0.01 in the full space (mean *±*s.e. across *n* = 31 sessions throughout; Fig. 6b). By comparison, decoding from five local PCs reached 0.64 *±*0.02, and matched random sub-spaces reached 0.58 *±*0.01. The communication subspace therefore captured lick-related activity more effectively than either high-variance local dimensions or arbitrary projections of the same dimensionality. Consistent with this preference, decoding from the communication-orthogonal complement fell to 0.56 *±* 0.01, close to chance.

This organization also depended on behavioral state. When the same active-block activity was projected onto communication axes estimated from passive replay, lick decoding remained above chance but was less accurate than with active axes (Fig. S7d). Together, these results show that much of the cortical activity useful for linear lick decoding lay within the dimensions that covaried with midbrain during engagement.

We next asked whether lick-related activity was similarly concentrated in the corresponding midbrain communication dimensions. In SCm/MRN, full-population decoding reached 0.85 *±*0.01, and the five active communication dimensions retained much of this performance, at 0.82 *±*0.02. However, local PCs performed similarly, at 0.81 *±*0.02, while matched random subspaces and the communication-orthogonal complement also supported substantial decoding (0.71 *±* 0.01 and 0.72 *±* 0.02, respectively; Figures 6b to 6c).

The cortex–midbrain contrast therefore concerned how lick-related activity was distributed across population dimensions. In VISm, decoding depended strongly on the communication subspace. In SCm/MRN, communication dimensions supported efficient low-dimensional decoding, but substantial lick information remained accessible outside them. Passive communication axes likewise supported midbrain lick decoding, although less accurately than active axes (Fig. S7d). The VISpl→SCm/MRN analysis reproduced this qualitative contrast between preferential cortical decoding within communication dimensions and broader midbrain decodability (Fig. S7d).

The preference for cortical communication dimensions could reflect a general concentration of task-related information or a more specific relationship to licking. To distinguish these possibilities, we applied the same subspace-decoding framework to eight-way image identity. Along VISm→SCm/MRN, image identity was strongly decodable from cortical activity, but its distribution differed from that of lick-related information: local PCs outperformed the active communication subspace, and the communication-orthogonal complement nearly recovered full-space decoding (Fig. S7e). Image decoding was comparatively weak across SCm/MRN subspaces. Thus, the active cortical communication dimensions preferentially captured lick-related activity without showing the same preference for image identity.

We then asked whether communication dimensions captured visual information more effectively at an earlier circuit stage. Along LGd→VISpl, communication subspaces supported substantially stronger image decoding than matched random subspaces in both source and target populations (Fig. S7f). The information accessible within communication dimensions therefore depended on the pathway: thalamocortical communication subspaces captured image information, whereas active cortical communication dimensions along VISm→SCm/MRN preferentially captured lick-related activity.

Together, these findings give behavioral meaning to the engagement-dependent covariance structure identified above. The active VISm communication subspace captured lick-related activity that was more broadly decodable in mid-brain, while image information showed a different distribution across subspaces and pathways. This links cross-area covariance geometry to the content of population activity, although decoding alone does not establish causal transmission between areas.

## Discussion

### Summary and conceptual advance

Our results support a circuit-level view of behavioral engagement in which population geometry and the inter-area modes through which activity is shared are jointly reorganized. Using large-scale Neuropixels recordings from the mouse thalamocortical– midbrain visual hierarchy, we found that active task engagement and passive replay define distinct neural population operating points. Relative to passive replay of matched visual stimuli, active engagement reduced mean activity, response participation, and dimensionality across multiple stages of the circuit. These findings extend prevailing accounts of state modulation, which have often emphasized local changes in gain, firing-rate baseline, recurrent amplification, or variability (19, 29, 30, 36). They instead point to a circuit-level view in which engagement-dependent population geometry reflects not only local dynamics, but also the structure of afferent drive from upstream populations.

### Input geometry as a model for active routing

The theory developed here provides a compact framework for active routing across brain states. In this framework, a target population is shaped by intrinsic fluctuations and recurrent dynamics, but also by the covariance geometry of its afferent drive. Structured source activity enters through an effective afferent coupling matrix and is filtered by target recurrent dynamics, so that the variance and dimensionality of source activity can constrain downstream dimensionality through separable mechanisms. This perspective reframes engagement-dependent modulation: behavioral state may reroute activity by changing which source modes are expressed, how much variance they carry, and how effectively they are coupled into downstream populations. The empirical source–target relationships were stage dependent: source and target activity were associated across all three channels, whereas dimensionality covaried across the LGd→VISpl and VISpl→VISm stages but not detectably at VISm→SCm/MRN.

### Cortex-to-midbrain communication geometry

The absence of a significant VISm–SCm/MRN dimensionality relationship marked a stage-specific limit of our minimal model and motivated a direct analysis of cross-area covariance geometry. During engagement, VISm–SCm/MRN shared covariance was concentrated into fewer effective modes, and the leading cross-area SVD subspaces showed comparatively weaker active–passive alignment than the dominant local PC subspaces. Thus, the cortex-to-midbrain transition involved a state-dependent reorganization of the dimensions along which VISm and SCm/MRN activity covaried, rather than a direct correspondence in population dimensionality.

### Different concentration of lick information in cortex and mid-brain

Subspace decoding linked engagement-dependent communication geometry to licking while revealing a cortex– midbrain asymmetry. In VISm, five engaged communication dimensions nearly recovered full-space decoding and outperformed matched local-PC and random subspaces, whereas decoding from the orthogonal complement approached chance level accuracy; the same pattern was reproduced in VISpl. In SCm/MRN, by contrast, communication and local-PC subspaces decoded similarly well, while random directions and the communication-orthogonal complement also remained informative, indicating that lick decodability was more broadly distributed across the mid-brain population space. Supplementary image decoding showed that this communication geometry supported different content across pathways: image identity was efficiently decoded from the LGd→VISpl communication subspace, but not preferentially from the engaged VIS→SCm/MRN communication subspace, and remained weakly decodable in SCm/MRN. Together, these results indicate pathway-specific decoding content within inter-area communication subspaces.

### Relation to gain, arousal, and local state modulation

These findings do not contradict classical accounts of gain modulation, locomotion-related modulation, or arousal-dependent changes in visual cortex (3, 4, 37–39). Rather, they suggest that such mechanisms are insufficient to explain the full population geometry observed here. Our analyses retained absolute population responses rather than subtracting a pre-stimulus baseline (1), thereby quantifying condition-specific operating points. Active and passive conditions can therefore differ in their full-state geometry even under matched sensory stimulation.

### Limitations and future directions

The main limitations of this study also point to the next experimental steps. Our source–target analyses follow the known thalamocortical– midbrain hierarchy but remain correlational; establishing causality will require pathway-specific perturbations. Engagement is also a composite state, and because active viewing preceded passive replay, the comparison is confounded by block order and elapsed time. Although within-block analyses argue against gradual drift, they cannot separate task context from reward availability, lick-spout presence, or other time-dependent factors. Future experiments combining large-scale recordings with behavioral, neuromodulatory, and pathway-specific measurements will be needed to disentangle these contributions.

The theory is likewise deliberately minimal: effective afferent couplings were modeled as independent zero-mean Gaussian variables with common variance *V*_*A*_ and without specificity to any particular modes. Indeed, biological connectivity may instead selectively couple particular source modes to target-population directions (34, 40), potentially contributing to the stage-specific transformations observed here. Recent work on networks with spatial organization in recurrent and afferent connectivity for example showed that interareal communication does not simply depend on the dimensionality of individual populations, but also on the alignment of within-population shared fluctuations across connected areas (41). Extending the framework to such more structured afferent connectivity will therefore be important. More broadly, input geometry provides a way to think about flexible routing in distributed circuits: behavioral state may shape computation by selecting which source modes are transmitted, amplified, or suppressed.

## Methods

### Dataset and analysis cohort

We analyzed first-day recording Familiar sessions from the Allen Visual Behavior Neuropixels dataset (1), comprising 48 sessions: 38 with image set G and 10 with image set H (Fig. S1b). Mice were trained on one of two sets of eight natural images until they met handoff criteria: peak d-prime *>* 1 for three consecutive sessions, at least 100 contingent non-aborted trials, and a mean of 120 rewards over the final three training sessions. During the active block, mice performed the rewarded visual change-detection task; during the passive block, the same image sequence was replayed with the lick spout retracted and no reward (Fig. S1a). Each image change was followed by a 3-s grace period in which no additional change could occur. Pre-mature licks aborted and restarted the trial, thereby delaying reward. Stimulus omissions occurred on 5% of non-change presentations. On the first recording day, mice viewed the trained Familiar image set; on the second day, they viewed a Novel set containing two Familiar images. Only first-day Familiar sessions were included.

### Spike counts, analysis windows, and area definitions

Units were retained if they had signal-to-noise ratio *>* 1, inter-spike-interval violations *<* 1, and firing rate *>* 0.1 Hz. Spikes were counted in 50-ms bins. Population-geometry analyses used bins from 0.10 to 0.25 s after image onset, corresponding to the final 150 ms of each 250-ms image presentation and the more stable sustained response (Fig. S1a). Omitted, novel, and image-change presentations were excluded; image changes elicited enhanced activity and therefore represented a distinct response regime from the repeated-image activity analyzed here.

Grouped circuit populations were defined from Allen structure acronyms as follows: LGd; primary and lateral visual cortex (VISpl) contained {VISp, VISrl, VISl, VISal}; medial visual cortex (VISm) contained {VISpm, VISam}; and motor-related superior colliculus and midbrain reticular nucleus (SCm/MRN) contained {SCig, SCiw, MRN}. Analyses required at least *N* = 30 quality-filtered neurons in each relevant population. Single-population analyses retained *n* = 16 sessions for LGd, *n* = 48 for VISpl, *n* = 47 for VISm, and *n* = 31 for SCm/MRN. Analyses of individual visual cortical areas retained *n* = 46 sessions for VISp, *n* = 44 for VISrl, *n* = 44 for VISl, *n* = 47 for VISal, *n* = 47 for VISpm, and *n* = 44 for VISam. Paired analyses retained *n* = 16 sessions for LGd→VISpl, *n* = 47 for VISpl→VISm, and *n* = 31 for VISm→SCm/MRN; VISpl→SCm/MRN analyses also retained *n* = 31 sessions.

### Active-block engagement analysis

Lick rate was computed in 1-s bins and smoothed with a 60-s window. Intervals were classified as unengaged when the smoothed lick rate remained below 20% of the session maximum for at least 90 s. Engaged–unengaged comparisons included the 12 sessions with at least 12 min of unengaged activity. After applying the *N* ≥ 30 criterion, this yielded *n* = 12 VISp, *n* = 10 VISrl, *n* = 10 VISl, *n* = 12 VISal, *n* = 12 VISpm, and *n* = 11 VISam sessions.

### Population statistics, PCA, and cross-area SVD

Scalar population statistics were computed from consecutive chunks of 200 viewing bins within each session, engagement state, and image. Mean activity was the average firing rate across neurons. Response participation was *D*(***r***) = E[***r***]^2^*/*E[***r***^2^], and dimensionality was *D*(***C***) = E[***λ***]^2^*/*E[***λ***^2^], where *λ*_*i*_ are the eigenvalues of the covariance matrix ***C***. Fano-factor participation was calculated analogously across neurons. For ratio analyses, chunk-level values were averaged within session, image, and engagement state, yielding one observation per session *×* image (8 images). Source–target regressions retained the chunk-level observations.

PCA and cross-area SVD were computed separately from the full set of retained 50-ms bins in each session *×* image *×* engagement-state group, rather than from the 200-bin chunks. We formed a samples-by-neurons spike-count matrix for each group and mean-centered each neuron without variance normalization. The local spike-count covariance was

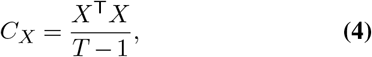

where *X* is the mean-centered activity matrix and *T* is the number of samples. Local population modes were the eigenvectors of *C*_*X*_ obtained by principal component analysis (PCA). For simultaneously recorded source and target activity matrices with matched time bins, the cross-area covariance was

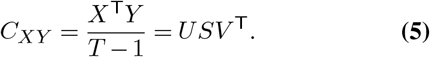

The leading columns of *U* and *V* define the source- and target-side communication subspaces, respectively; together, these paired SVD subspaces constitute the cross-area communication channel. Mode-similarity and subspace-capture analyses used the leading *k* = 5 modes; decoding analyses also used *k* = 5.

### Source–target regression

For LGd→VISpl, VISpl→VISm, and VISm→SCm/MRN, multiple linear regressions were fit separately within each session and active/passive block. Each observation comprised matched source and target statistics from one image *×* 200-bin chunk. Target mean activity or dimensionality was predicted from running speed, pupil area, source mean activity, and source dimensionality. Predictors and outcomes were z-scored within each session and condition, placing all coefficients on a common scale. Example source–target plots (Fig. 4a) show these chunk-level observations with ordinary least-squares fits and 95% confidence intervals. Group-level inference was performed on the resulting session-level coefficients.

### Mode and subspace similarity across states

For mode-wise active–passive similarity, we calculated absolute pair-wise cosine similarities between the leading five active and passive modes. The Hungarian algorithm identified the one-to-one pairing that maximized total cosine similarity while allowing mode order to differ between conditions. Similarities and directional alignment indices were computed separately for each session and image (8 images; *n* = 31 simultaneously recorded VISm–SCm/MRN and VISpl–SCm/MRN sessions).

Directional subspace capture followed the variance-capture alignment index introduced in (35). If *P*^*A*^ and *P*^*P*^ are the leading *k* = 5 active and passive local-PC bases, respectively, the fraction of passive covariance captured by the active subspace was

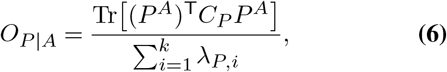

where *C*_*P*_ is the passive local covariance matrix and *λ*_*P,i*_ are its leading eigenvalues. The reverse direction, *O*_*A*|*P*_ , was obtained by exchanging active and passive quantities.

We generalized the same alignment principle to cross-area dynamics by asking how much cross-covariance from one condition was captured jointly by the paired source and target SVD subspaces defined in the other condition. For passive cross-covariance captured by the active communication subspaces,

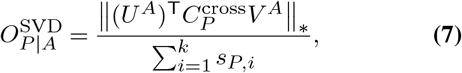

where *U*^*A*^ and *V* ^*A*^ contain the leading *k* = 5 active source and target singular vectors, *s*_*P,i*_ are the leading passive singular values, and ∥·∥_∗_ denotes the nuclear norm. This quantity measures the singular-value mass of the passive cross-covariance retained after projection onto the active communication subspaces. The reverse direction was obtained by exchanging active and passive quantities.

### Lick decoding and subspace readouts

Lick decoding used active-block activity only. Each sample was a 50-ms bin from the 0.10–0.25 s response window of a non-change presentation and was labeled according to whether a lick occurred in that bin. Change and omitted presentations were excluded to match the repeated-image regime used for the population-geometry analyses. Rewarded post-change licks typically occurred during the inter-stimulus gray period; lick-positive samples in the analyzed window therefore represented non-rewarded error licks rather than correct change-detection responses. Spike counts were mean-centered but not variance-normalized. Sessions were required to contain at least 30 lick-positive samples; all sessions eligible for the paired VISm→SCm/MRN and VISpl→SCm/MRN analyses met this criterion.

We used *L*_2_-regularized logistic regression with inverse regularization strength *C* = 0.1. All lick-positive samples were retained, and no-lick samples were randomly undersampled to balance the classes. Undersampling was repeated ten times; each repeat used five-fold stratified cross-validation, and performance was averaged across repeats within each session. Decoding performance was quantified by balanced accuracy,

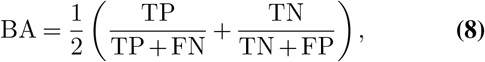

the mean of sensitivity and specificity. Because evaluation samples were balanced by undersampling, balanced accuracy was numerically equivalent to accuracy.

Subspace decoding used *k* = 5 and compared (i) the full population space; (ii) the active communication subspace from cross-area SVD; (iii) the active local-PC subspace; (iv) random orthonormal five-dimensional subspaces; and (v) the orthogonal complement of the active communication sub-space, computed as *X* − *XBB*^T^ for basis *B*. Five random subspaces were sampled per undersampling repeat. Supplementary controls projected active activity onto communication and PCA axes estimated from passive viewing; passive activity itself was not decoded.

### Supplementary image-identity decoding

Image decoding was an eight-class supplementary analysis restricted to image set G. Each sample was one image presentation, with spike counts averaged over 0.10–0.25 s after image onset. Presentations were included irrespective of lick status, which was not used as a label or selection criterion. SVD and PCA bases were computed from population matrices (image presentations *×* neurons) separately for active and passive blocks, and all subspace readouts used *k* = 5. Image identities were decoded in the full population space, active communication subspace, active local-PC subspace, random five-dimensional subspaces, and the orthogonal complement of the active communication subspace. Additional controls projected active image responses onto passive communication or passive PCA axes; passive activity itself was not decoded. Five random orthonormal subspaces were sampled per session.

The decoder was *L*_2_-regularized logistic regression with *C* = 0.1 and five-fold stratified cross-validation. Multiclass balanced accuracy was the macro-average recall,

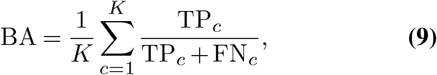

with *K* = 8 and chance level 1*/*8. After applying these criteria, the VISm→SCm/MRN and VISpl→SCm/MRN analyses each included *n* = 26 sessions, whereas LGd→VISpl included *n* = 13 sessions.

### Statistical analysis and figure conventions

All reported *t*-tests were two-sided. Metric ratios were tested against the null value of 1 with one-sample *t*-tests at the observation level specified in each caption. Session-level regression coefficients were tested against zero with one-sample *t*-tests, and active and passive coefficients from matched sessions were compared with paired *t*-tests. Reported *p* values were not adjusted for multiple comparisons. Unless otherwise noted, boxplots show the median, interquartile range, and whiskers extending to 1.5*×* the interquartile range. Violin plots show the observation distribution with an inner box of the same form as the boxplots. Means, SEMs, and confidence intervals are identified explicitly in the corresponding captions.

## Acknowledgments

This work was partially funded by the Deutsche Forschungsgemeinschaft (DFG, German Research Foundation) as part of the SPP 2205-533396241. Loris Amalberti is grateful to the Azrieli Foundation for the award of an Azrieli Fellowship. We thank the Allen Institute for Brain Science founder, Paul G. Allen, for his vision, encouragement, and support.

---

**Fig. S1.**
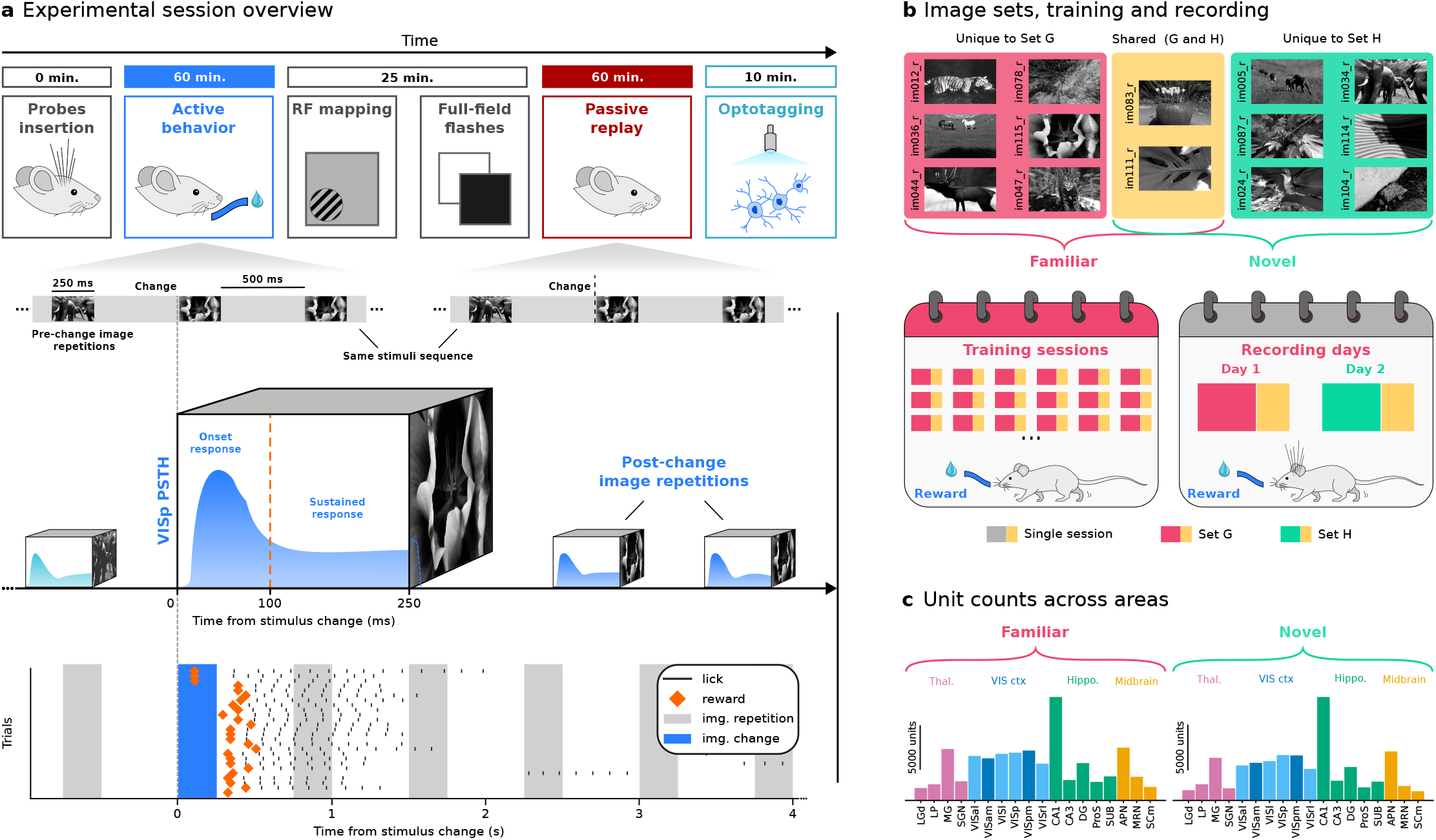
Recording-day structure, image sets, and unit yields. a) Neuropixels recording-day sequence: visual change-detection task (∼60 min), receptive-field mapping and full-field flashes (∼25 min), passive replay with the lick spout retracted and no reward (∼60 min), and optotagging (∼10 min). Active and passive blocks used the same image sequence. The schematic VISp peri-stimulus time histogram marks the onset-response (20–100 ms (1)) and sustained-response (100–250 ms) windows; analyses used the sustained response. Change-aligned lick and reward rasters are shown for an example session. b) Natural-image sets used during training and recording. Mice viewed the trained eight-image set (Familiar) on the first recording day and a Novel eight-image set, containing two Familiar images, on the second day. c) Pooled numbers of quality-filtered units (cf. Methods) by area across Familiar sessions, grouped by thalamus, visual cortex, hippocampus, and midbrain; yields from Novel sessions are shown for comparison.

**Fig. S2.**
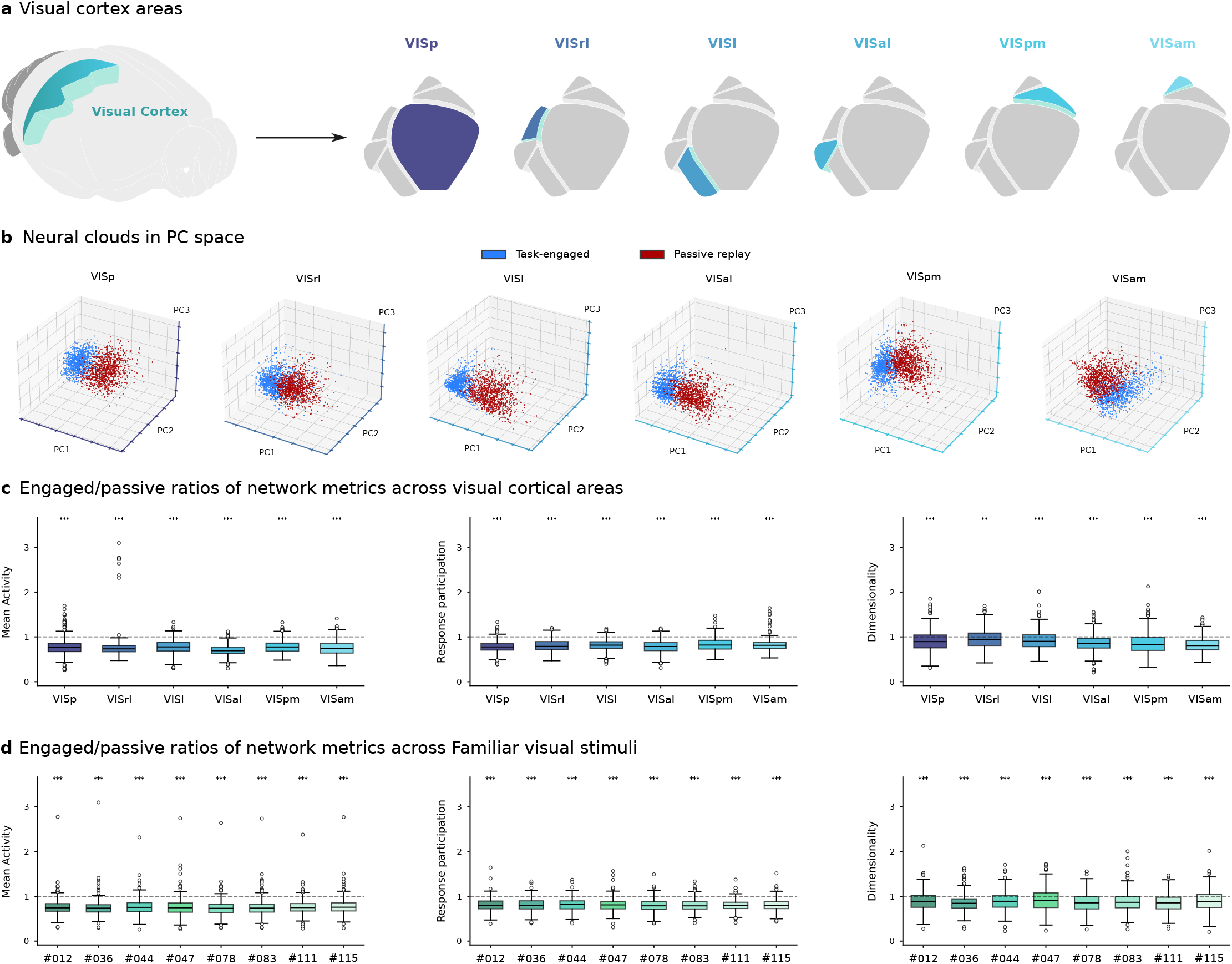
Engagement shifts population operating points and compresses geometry across visual cortex. a) Anatomical localization of the recorded visual cortical areas: primary visual cortex (VISp), rostrolateral (VISrl), lateral (VISl), anterolateral (VISal), posteromedial (VISpm), and anteromedial (VISam) visual cortex. b) Low-dimensional PCA projections of population responses across visual cortical areas during active engagement and passive viewing. Active response clouds are shown in blue and passive response clouds in red. Their separation illustrates the engagement-dependent shift in population operating points across visual cortex. c) Task-engaged/passive ratios of mean activity, response participation, and dimensionality across VISp and lateral (VISrl, VISl, VISal) and medial (VISpm, VISam) visual cortical areas. Boxplots show session *×* image observations (8 images; *n* = 46 VISp, 44 VISrl, 44 VISl, 47 VISal, 47 VISpm, and 44 VISam sessions). The dashed line marks a ratio of 1. Asterisks denote two-sided one-sample *t*-tests against 1: ^∗∗^*p <* 0.01, ^∗∗∗^*p <* 0.001. d) Task-engaged/passive ratios of the same metrics for each Familiar stimulus in image set G. Boxplots show session *×* visual-area observations for each of the 8 images (*n* = 38 sessions overall; *n* = 37 VISp, 34 VISrl, 35 VISl, 37 VISal, 38 VISpm, and 36 VISam sessions). The dashed line marks a ratio of 1. Asterisks denote two-sided one-sample *t*-tests against 1: ^∗∗∗^*p <* 0.001.

**Fig. S3.**
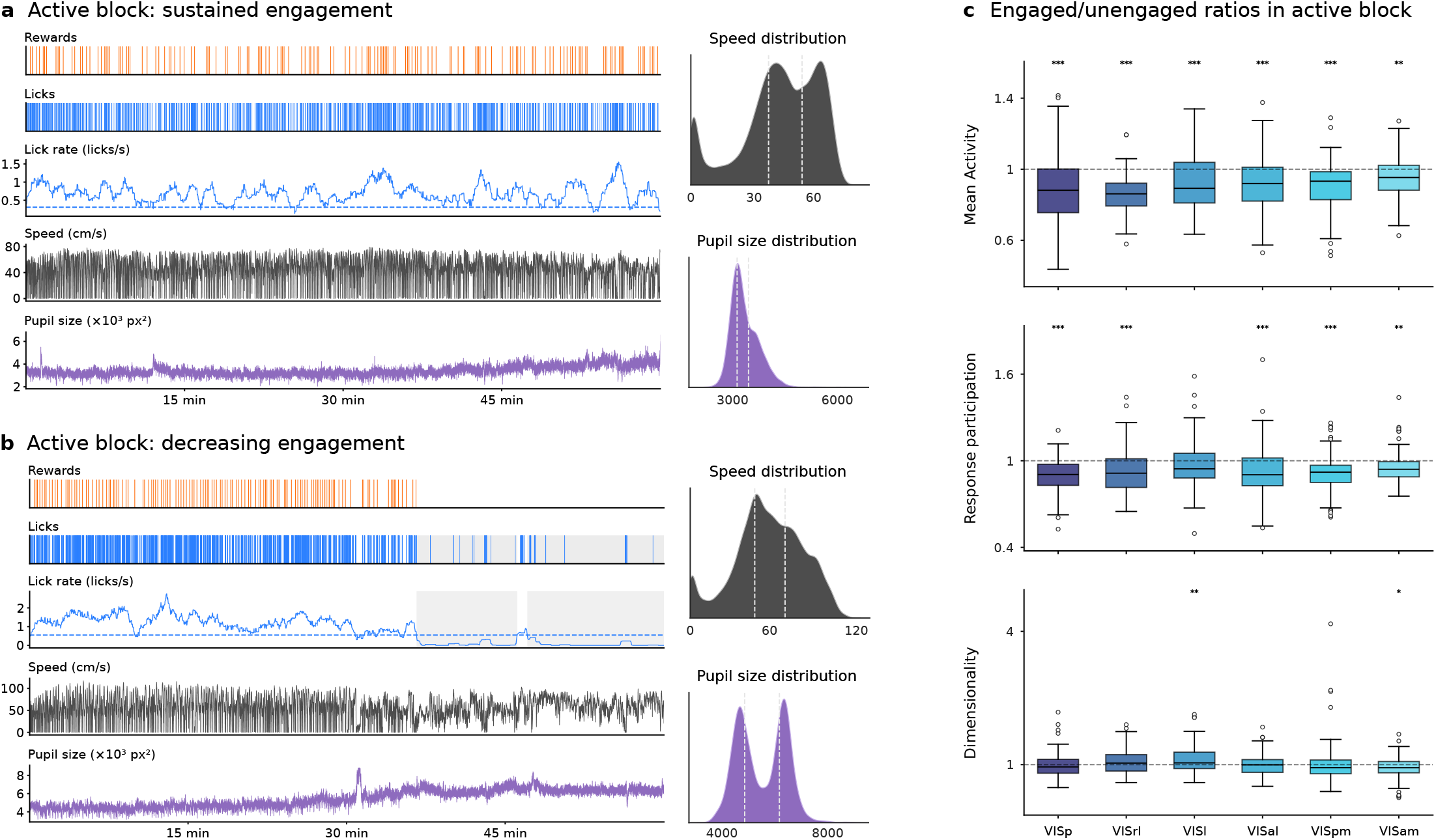
Within-block loss of engagement and associated population metrics. a) Behavioral traces from an example session with sustained engagement throughout the active block. Running-speed and pupil-size distributions are shown with tercile boundaries. b) Example session in which engagement declined, identified from licking and lick-rate dynamics (shading). Running-speed and pupil-size distributions are shown as in a. c) Engaged/unengaged ratios of mean activity, response participation, and dimensionality during within-block loss of engagement. Boxplots show session *×* image observations (8 images; *n* = 12 sessions overall; *n* = 12 VISp, 10 VISrl, 10 VISl, 12 VISal, 12 VISpm, and 11 VISam sessions). The dashed line marks a ratio of 1. Asterisks denote two-sided one-sample *t*-tests against 1: ^∗^*p <* 0.05, ^∗∗^*p <* 0.01, ^∗∗∗^*p <* 0.001.

**Fig. S4.**
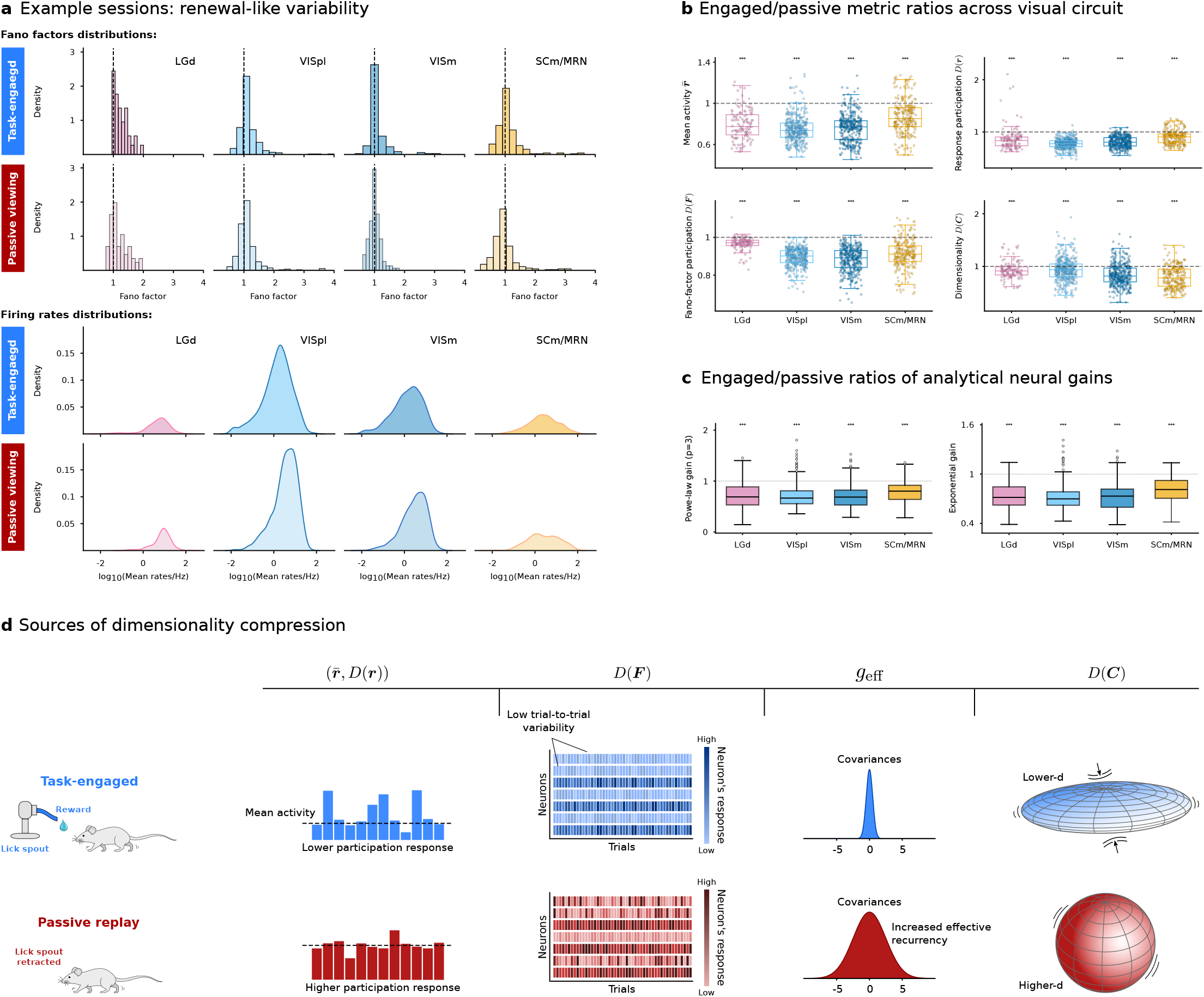
Engagement-dependent changes in local population statistics and analytical neural gains across the visual circuit. a) Example-session distributions of single-neuron Fano factors and firing rates across LGd, VISpl, VISm, and SCm/MRN during task engagement and passive viewing. Dashed lines in the Fano-factor distributions mark the Poisson expectation, for which spike-count variance equals the mean. b) Task-engaged/passive ratios of mean activity, response participation, Fano-factor participation, and dimensionality across LGd, VISpl, VISm, and SCm/MRN. Boxplots and overlaid points show session *×* image observations (8 images; *n* = 16 LGd, 48 VISpl, 47 VISm, and 31 SCm/MRN sessions); the boxplot-only summary is shown in Fig. 2f. The dashed line marks a ratio of 1. Asterisks denote two-sided one-sample *t*-tests against 1: ^∗∗∗^*p <* 0.001. c) Task-engaged/passive ratios of analytically estimated neuronal gain moments for the nonlinear transfer functions considered in the local theory. Boxplots show session *×* image observations as in b. The dashed line marks a ratio of 1. Asterisks denote two-sided one-sample *t*-tests against 1: ^∗∗∗^*p <* 0.001. d) Conceptual summary of local population quantities associated with engagement-dependent dimensionality compression. Relative to passive viewing, task engagement is characterized by lower mean activity, response participation, Fano-factor participation, analytically estimated neuronal gain/recurrency (19, 24), and dimensionality.

**Fig. S5.**
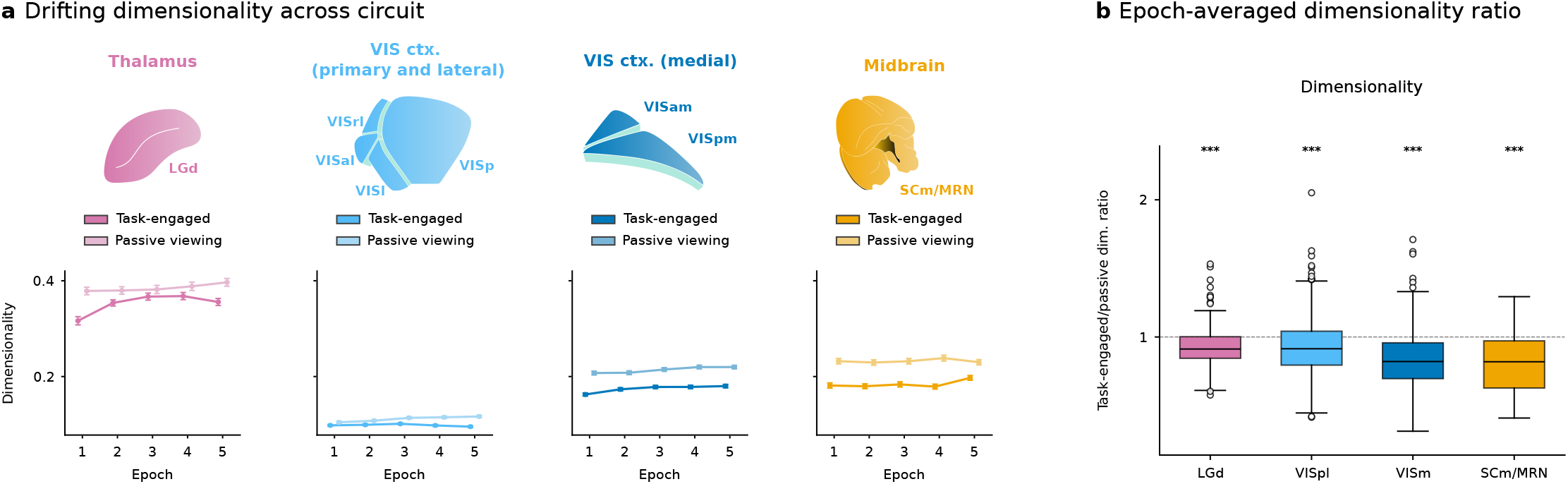
Engagement-dependent dimensionality compression is robust to population size and within-block time. a) Dimensionality across successive 11-min epochs after discarding the first 5 min of active and passive blocks. Curves and shading show mean *±* SEM across session *×* image observations (*n* = 16 LGd, 48 VISpl, 47 VISm, and 31 SCm/MRN sessions; 8 images). b) Task-engaged/passive ratio of dimensionality averaged across within-block epochs. Boxplots show session *×* image observations as in a. The dashed line marks a ratio of 1. Asterisks denote two-sided one-sample *t*-tests against 1: ^∗∗∗^*p <* 0.001.

**Fig. S6.**
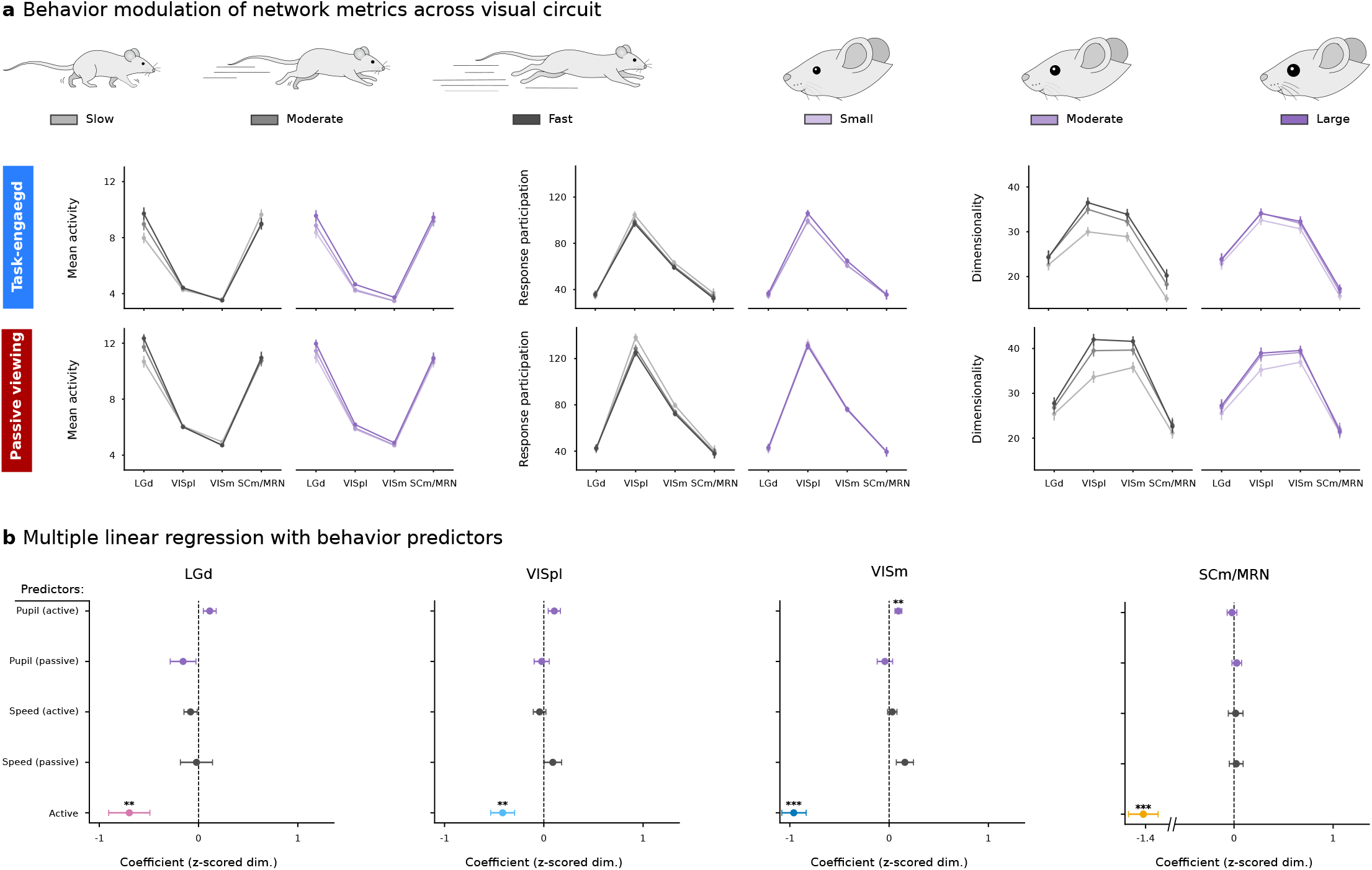
Locomotion and arousal do not account for engagement-dependent dimensionality. a) Mean activity, response participation (shown as the effective number of participating neurons, *ND*(***r***)), and dimensionality in LGd, VISpl, VISm, and SCm/MRN, grouped by running-speed tercile (slow, moderate, fast) or pupil-area tercile (small, moderate, large) within active and passive blocks. Points and error bars show means and 95% confidence intervals across session *×* image observations (8 images; *n* = 16 LGd, 48 VISpl, 47 VISm, and 31 SCm/MRN sessions). b) Standardized coefficients from models predicting dimensionality from active/passive condition and condition-specific running-speed and pupil-area terms. Points and error bars show mean *±* SEM across session-level coefficients (session numbers as in a). Asterisks denote two-sided one-sample *t*-tests against zero: ^∗∗^*p <* 0.01, ^∗∗∗^*p <* 0.001.

**Fig. S7.**
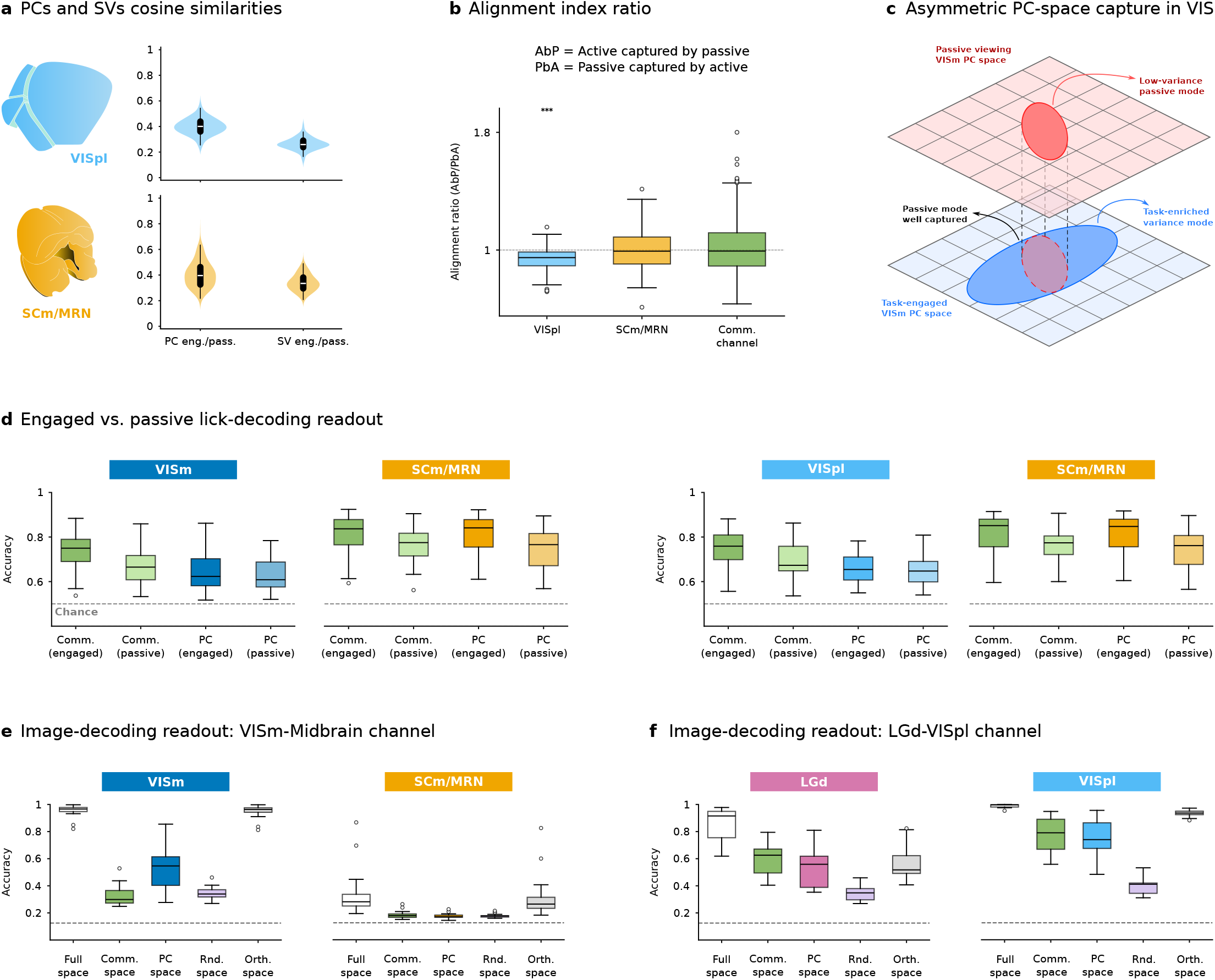
State-dependent subspace alignment and decoding controls across visual pathways. a) Mean absolute cosine similarity between task-engaged and passive modes for the leading *k* = 5 local principal components and cross-area singular vectors. Modes were paired as in Fig. 5d. Violin plots show session *×* image observations (*n* = 31 sessions, 8 images). b) Ratio of directional cross-condition capture (AbP/PbA) for the leading *k* = 5 local principal-component and communication subspaces. Boxplots show session *×* image observations as in a; the dashed line marks a ratio of 1. Asterisks denote two-sided one-sample *t*-tests against 1: ^∗∗∗^*p <* 0.001. c) Schematic of asymmetric PC-subspace capture in visual cortex. The task-engaged subspace captures a passive mode while additionally recruiting a task-enriched high-variance direction, yielding stronger passive-by-active than active-by-passive capture. d) Five-fold cross-validated lick decoding from active-block activity projected onto task-engaged or passive communication and principal-component axes for the VISm–SCm/MRN and VISpl–SCm/MRN channels. Boxplots show session-level means (*n* = 31 sessions); the dashed line marks chance (0.5). e) Eight-way image-identity decoding along the VISm–SCm/MRN channel (*n* = 26 sessions, image set G), comparing the full population space, the communication subspace, local principal components, a random matched-dimensional subspace, and the orthogonal complement of the communication modes. f) Corresponding image-identity decoding along the LGd–VISpl channel (*n* = 13 sessions, image set G). In e and f, boxplots show session-level balanced accuracy and chance is 0.125.

