## Supplementary Material for "Task Engagement Gates Interareal Communication Geometry in the Mouse Thalamocortical–Midbrain Visual Circuit"

#### Overview

This Supplementary Information provides the theoretical derivation supporting the input-geometry framework introduced in the main text. We define a rate-based recurrent network with structured afferent drive, linearize around a stable operating point, and characterize the geometry of source-area fluctuations. A path-integral calculation, extending the purely local formalism of Layer, Helias, and Dahmen [1] towards correlated external inputs, yields the target covariance moments and dimensionality. We then examine limiting cases without afferent drive and extend the nonlinear rate formulation to renewal-like spiking variability, recovering and generalizing the result of Tian et al. [2].

#### Contents

**1 Nonlinear rate model and linearization** 1

**2 Linear model with structured afferent input** 3

**3 Source covariance geometry and moment matching** 5

**4 Path-integral formulation and target dimensionality** 6

**5 Limiting cases and Poisson-like variability** 10

#### 1 Nonlinear rate model and linearization

We consider a target population of $N$ interacting neurons whose activity is described by their firing rates $r_i(t)$. Each target neuron $i$ is driven by three contributions: recurrent input from other neurons in the same target population, structured afferent input from $M$ neurons of a source population with firing rates $s_\alpha(t)$, with $\alpha \in \{1, \ldots, M\}$, and intrinsic fluctuations $\xi_i(t)$. The target network dynamics is therefore governed by $N$ coupled stochastic rate equations:

$$\tau \dot{r}_i(t) = -r_i(t) + f\left(\sum_{j=1}^{N} W_{ij}\, r_j(t - d_1) + \sum_{\alpha=1}^{M} \Lambda_{i\alpha}\, s_\alpha(t - d_2)\right) + \xi_i(t)\,. \tag{1}$$

Here, $\tau$ is the relaxation time constant of the target firing-rate dynamics, $f(\cdot)$ is an element-wise nonlinear transfer function, $W_{ij}$ is the recurrent connectivity from neuron $j$ to neuron $i$, $\Lambda_{i\alpha}$ is the afferent coupling from source neuron $\alpha$ to target neuron $i$, and $d_1$ and $d_2$ are transmission delays.

Following previous recurrent random network models [3, 4, 5], we here, for simplicity, take the target connectivity to be independently drawn from a zero-mean Gaussian ensemble and model intrinsic fluctuations as independent Gaussian noise

$$W_{ij} \sim \mathcal{N}\left(0, \frac{g^2}{N}\right), \qquad \xi_i(t) \sim \mathcal{N}(0, D_{0,i}), \tag{2}$$

where $D_{0,i}$ is the intrinsic noise strength of target neuron $i$. The parameter $g$ controls the spectral radius of the connectivity matrix, and, in the absence of gain heterogeneity and afferent drive, the condition $g < 1$ ensures stability of the random recurrent network [3, 6]. Note that generalizations to more complicated local connectivities, such as sparse couplings between multiple populations, could be derived analogously to previously published formalisms [1, 7].

We assume the system operates around a stable operating point, where mean rates of the deterministic equation satisfy

$$r_i^* = f(h_i^*), \qquad \text{with: } h_i^* = \sum_j W_{ij}\, r_j^* + \sum_\alpha \Lambda_{i\alpha}\, s_\alpha^*, \tag{3}$$

where $h_i^*$ is the total steady-state synaptic input to target neuron $i$.
To study the response to small perturbations, we linearize the dynamics around this fixed point. Writing $r_i(t) = r_i^* + \delta r_i(t)$ and $s_\alpha(t) = s_\alpha^* + \delta s_\alpha(t)$, the total input current expands as

$$h_i(t) = h_i^* + \sum_{j=1}^{N} W_{ij}\, \delta r_j(t - d_1) + \sum_{\alpha=1}^{M} \Lambda_{i\alpha}\, \delta s_\alpha(t - d_2). \tag{4}$$

We then ask how recurrent and afferent perturbations propagate through the target network around this fixed operating point. A first-order Taylor expansion of the nonlinearity around $h_i^*$ gives

$$f(h_i^* + \delta h_i) \simeq f(h_i^*) + f'(h_i^*)\, \delta h_i + \mathcal{O}(\delta h_i^2), \tag{5}$$

where $\delta h_i = \sum_j W_{ij}\, \delta r_j(t - d_1) + \sum_\alpha \Lambda_{i\alpha}\, \delta s_\alpha(t - d_2)$ and we use the shorthand notation $f_i^* \equiv f'(h_i^*) \equiv \partial_i f(h_i^*)$. In the expression above, the derivative $f_i^*$ is the local gain of target neuron $i$ at the operating point, denoting how strongly small changes in recurrent and afferent perturbations are converted into changes in firing rate.
Substituting the first-order expansion into Eq. (1) and using the fixed-point condition in Eq. (3), the deterministic terms cancel, and the linearized dynamics for the rate fluctuations become

$$\tau \dot{\delta r}_i(t) = -\delta r_i(t) + \sum_j f_i^* W_{ij}\, \delta r_j(t - d_1) + \sum_\alpha f_i^* \Lambda_{i\alpha}\, \delta s_\alpha(t - d_2) + \xi_i(t). \tag{6}$$

Thus, the nonlinearity affects the fluctuation dynamics only through the local slope $f_i^*$, which can be collected into a diagonal matrix

$$\begin{bmatrix} f_1^* & 0 & \cdots & 0 \\ 0 & \ddots & & \vdots \\ \vdots & & \ddots & 0 \\ 0 & \cdots & 0 & f_N^* \end{bmatrix} = \mathrm{diag}(f_1^*, \ldots, f_N^*) . \tag{7}$$

This matrix notation makes explicit that gain renormalization acts row-wise on incoming synaptic weights: all inputs to the same postsynaptic neuron are multiplied by the same local response gain.

We therefore define the *effective* connectivity $W^*_{ij} \equiv f^*_i\, W_{ij}$, and, similarly, the gain-rescaled afferent coupling $\Lambda^*_{i\alpha} \equiv f^*_i \Lambda_{i\alpha}$.
We next characterize the statistics of the effective connectivity. Treating the gain factors $f^*_i$ and the bare synaptic weights $W_{ij}$ as independent to leading order in the large-$N$ limit[1], the first two cumulants of $W^*_{ij}$ are

$$\kappa^*_{1,ij} = \mathbb{E}\big[W^*_{ij}\big] = \mathbb{E}[W_{ij}]\,\mathbb{E}[f^*_i] = 0\,, \tag{8a}$$
$$\kappa^*_{2,ijkl} = \mathbb{E}\big[W^*_{ij}W^*_{k\ell}\big] - \mathbb{E}\big[W^*_{ij}\big]\,\mathbb{E}[W^*_{k\ell}] = \mathbb{E}[W_{ij}W_{k\ell}]\,\mathbb{E}[f^*_i f^*_k]\ . \tag{8b}$$

Because we assumed the bare connectivity to have zero mean and no correlation among distinct rows and columns, the only surviving second cumulant is

$$\kappa^*_{2,ijkl}\big|_{i=k,\,j=\ell} = \mathbb{E}\big[W^2_{ij}\big]\,\mathbb{E}\big[(f^*_i)^2\big] = \frac{g^2}{N}\,\mathbb{E}\big[(f^*_i)^2\big]\ . \tag{9}$$

The effect of gain heterogeneity on the synaptic weights is therefore captured, at this level of approximation, by a renormalization of the connectivity variance. Equivalently,

$$W_{ij} \sim \mathcal{N}\left(0, \frac{g^2}{N}\right) \implies W^*_{ij} \sim \mathcal{N}\left(0, \frac{g^2_{\text{eff}}}{N}\right)\,, \qquad g^2_{\text{eff}} = g^2\,\mathbb{E}\big[(f^*_i)^2\big]\ . \tag{10}$$

where $g_{\text{eff}}$ is the effective spectral radius. Once again stability of the linearized network requires $g^2_{\text{eff}} < 1$, meaning $g < 1/\sqrt{\mathbb{E}[(f^*_i)^2]}$.
For compact notation, we now rename the target fluctuations $\delta r_i \equiv x_i$ and the source fluctuations $\delta s_\alpha \equiv y_\alpha$, yielding the effective linear model

$$\tau \dot{x}_i(t) = -x_i(t) + \sum_j W^*_{ij}\, x_j(t-d_1) + \sum_\alpha \Lambda^*_{i\alpha}\, y_\alpha(t-d_2) + \xi_i(t)\,. \tag{11}$$

#### 2 Linear model with structured afferent input

We now analyze the effective linear model introduced in Eq. (11). By construction source fluctuations are zero mean and we assume them to be temporally white,

$$\mathbb{E}[y_\alpha(t)] = 0, \qquad \operatorname{Cov}\big[y_\alpha(t), y_\beta(t')\big] = \Sigma_{s,\alpha\beta}\,\delta(t-t')\,, \tag{12}$$

where $\boldsymbol{\Sigma}_s$ is the covariance matrix of source-area fluctuations, capturing the second-order geometry of the afferent activity that is transmitted to the target population. To compute the covariance of target fluctuations, we work in the frequency domain. We consider the Fourier transforms

$$\tilde{x}_i(\omega) = \int_{-\infty}^{\infty} dt\, e^{-i\omega t}\, x_i(t)\,, \qquad \tilde{y}_\alpha(\omega) = \int_{-\infty}^{\infty} dt\, e^{-i\omega t}\, y_\alpha(t)\,, \tag{13}$$

and analogously define $\tilde{\xi}_i(\omega)$ for the intrinsic noise. Taking the Fourier transform of (11) yields

$$(i\omega\tau + 1)\,\tilde{x}_i(\omega) = e^{-i\omega d_1}\sum_{j=1}^{N} W^*_{ij}\,\tilde{x}_j(\omega) + e^{-i\omega d_2}\sum_{\alpha=1}^{M} \Lambda^*_{i\alpha}\,\tilde{y}_\alpha(\omega) + \tilde{\xi}_i(\omega)\,. \tag{14}$$

[1] Although the operating point satisfies $r^*_i = f\Big(\sum_j W_{ij} r^*_j + \sum_\alpha \Lambda_{i\alpha} s^*_\alpha\Big)$, its dependence on any individual entry $W_{ij}$ is weak when $N$ is large; we therefore treat $f^*_i$ and $W_{ij}$ as independent to leading order.

It is convenient to define a composite noise variable,

$$\tilde{\eta}_i(\omega) = e^{-i\omega d_2} \sum_{\alpha=1}^{M} \Lambda^*_{i\alpha}\, \tilde{y}_\alpha(\omega) + \tilde{\xi}_i(\omega)\,, \tag{15}$$

which collects the intrinsic and afferent noise sources. Because $\tilde{\boldsymbol{y}}$ and $\tilde{\boldsymbol{\xi}}$ are independent and Gaussian, the composite noise is zero mean with covariance

$$\mathbb{E}[\tilde{\boldsymbol{\eta}}(\omega)] = \mathbf{0}\,, \qquad \mathbb{E}\Big[\tilde{\boldsymbol{\eta}}(\omega)\, \tilde{\boldsymbol{\eta}}^\mathsf{T}(\omega^*)\Big] = 2\pi\, \delta(\omega+\omega^*)\, \Big(\boldsymbol{D}_0 + e^{-i\omega d_2} \boldsymbol{\Lambda}^*\, \boldsymbol{\Sigma}_s\, \boldsymbol{\Lambda}^{*\mathsf{T}} e^{-i\omega^* d_2}\Big)\,, \tag{16}$$

where $\boldsymbol{D}_0 = \mathrm{diag}(D_{0,1}, \ldots, D_{0,N})$.
Solving Eq. (14) for $\tilde{\boldsymbol{x}}(\omega)$ and computing the cross-spectral density gives

$$\begin{aligned}\mathbb{E}\Big[\tilde{\boldsymbol{x}}(\omega)\, \tilde{\boldsymbol{x}}^\mathsf{T}(\omega^*)\Big] &= 2\pi\, \delta(\omega+\omega^*)\, \tilde{\boldsymbol{C}}(\omega^*) \\ &= 2\pi\, \delta(\omega+\omega^*)\, \boldsymbol{G}(\omega)\, \Big(\boldsymbol{D}_0 + e^{-i\omega d_2} \boldsymbol{\Lambda}^*\, \boldsymbol{\Sigma}_s\, \boldsymbol{\Lambda}^{*\mathsf{T}} e^{-i\omega^* d_2}\Big)\, \boldsymbol{G}^\mathsf{T}(\omega^*)\,,\end{aligned} \tag{17}$$

where $\boldsymbol{G}(\omega) = \big[(i\omega\tau + 1)\,\mathbb{I} - e^{-i\omega d_1}\, \boldsymbol{W}^*\big]^{-1}$ is the resolvent of the linearized dynamics. This resolvent describes how recurrent interactions filter both intrinsic and afferent fluctuations before they appear in the target population.
Employing the Wiener–Khinchin theorem, the time-lag integrated covariance is given by the zero-frequency cross-spectrum. We use this zero-frequency covariance as the theoretical analogue of the experimentally estimated trial-to-trial covariance of population activity [4, 1]. At $\omega = 0$, delay phase factors become irrelevant ($e^{-i\omega d_1} = e^{-i\omega d_2} = 1$), and Eq. (17) reduces to

$$\boldsymbol{C} \equiv \tilde{\boldsymbol{C}}(0) = (\mathbb{I} - \boldsymbol{W}^*)^{-1}\, \Big(\boldsymbol{D}_0 + \boldsymbol{\Lambda}^*\, \boldsymbol{\Sigma}_s\, \boldsymbol{\Lambda}^{*\mathsf{T}}\Big)\, (\mathbb{I} - \boldsymbol{W}^*)^{-\mathsf{T}}\,. \tag{18}$$

This expression shows that target covariance is shaped jointly by effective recurrent amplification through $\boldsymbol{W}^*$, intrinsic variability $\boldsymbol{D}_0$, and the afferent-drive geometry encoded in $\boldsymbol{\Sigma}_s$ and $\boldsymbol{\Lambda}^*$.
Next, to isolate the contribution of individual source covariance modes, we diagonalize the source covariance matrix. Because $\boldsymbol{\Sigma}_s$ is symmetric and positive semidefinite, it admits the eigendecomposition

$$\boldsymbol{\Sigma}_s = \boldsymbol{P}\, \boldsymbol{D}_s\, \boldsymbol{P}^\mathsf{T}, \qquad \boldsymbol{D}_s = \mathrm{diag}(\lambda_1, \ldots, \lambda_M)\,, \tag{19}$$

where $\boldsymbol{P}$ contains the orthonormal source eigenvectors and $\boldsymbol{D}_s$ contains the corresponding eigenvalues $\lambda_\alpha$. Each eigenvalue $\lambda_\alpha$ is the variance carried by source mode $\alpha$, with large eigenvalues corresponding to dominant source fluctuation modes, while small eigenvalues corresponding to weakly expressed modes.
Substituting Eq. (19) into Eq. (18) and absorbing the source eigenvectors into the afferent coupling gives

$$\boldsymbol{C} = (\mathbb{I} - \boldsymbol{W}^*)^{-1}\, \Big(\boldsymbol{D}_0 + \boldsymbol{\Lambda}^\dagger\, \boldsymbol{D}_s\, \boldsymbol{\Lambda}^{\dagger\mathsf{T}}\Big)\, (\mathbb{I} - \boldsymbol{W}^*)^{-\mathsf{T}}\,, \tag{20}$$

where $\boldsymbol{\Lambda}^\dagger \equiv \boldsymbol{\Lambda}^*\, \boldsymbol{P}$ is the afferent coupling expressed in the source eigenmode basis. This rotation does not change the total source covariance, instead it rewrites the same source fluctuations in coordinates where different source modes are uncorrelated. In this basis, the contribution of each source mode to the target covariance can be read directly from its variance $\lambda_\alpha$ and coupling pattern $\boldsymbol{\Lambda}^\dagger$.
Equivalently, we may rotate the source fluctuations as $\boldsymbol{m}(t) \equiv \boldsymbol{P}^\mathsf{T}\, \boldsymbol{y}(t)$. Since Gaussianity is preserved under linear transformations, we have

$$\boldsymbol{y}(t) \sim \mathcal{N}(\mathbf{0}, \boldsymbol{\Sigma}_s) \quad \Longrightarrow \quad \boldsymbol{m}(t) \sim \mathcal{N}\Big(\mathbf{0}, \boldsymbol{P}^\mathsf{T} \boldsymbol{\Sigma}_s \boldsymbol{P}\Big) = \mathcal{N}(\mathbf{0}, \boldsymbol{D}_s)\,, \tag{21}$$

with independent mode variances $\lambda_\alpha$ along the diagonal of $\boldsymbol{D}_s$. The time-domain dynamics can therefore be written as

$$\tau \dot{x}_i(t) = -x_i(t) + \sum_j W^*_{ij}\, x_j(t-d_1) + \sum_\alpha \Lambda^\dagger_{i\alpha}\, m_\alpha(t-d_2) + \xi_i(t)\,. \tag{22}$$

In this representation, low-dimensional source activity corresponds to the case in which a small number of $\lambda_\alpha$ dominate the source spectrum, while high-dimensional source activity corresponds to variance distributed more evenly across many modes.

The rotated entries $\Lambda^\dagger_{i\alpha}$ can be positive or negative. We here model them as zero-mean Gaussian random variables

$$\Lambda^\dagger_{i\alpha} \sim \mathcal{N}(0, V)\,, \tag{23}$$

which provides a tractable first approximation to the distribution of effective afferent coupling coefficients in the source-covariance eigenbasis. For notational simplicity, in the main text we suppress the asterisks and daggers and write $\boldsymbol{f}^* \to \boldsymbol{f}$, $\boldsymbol{W}^* \to \boldsymbol{W}$ and $\boldsymbol{\Lambda}^\dagger \to \boldsymbol{A}$; these quantities are nevertheless understood as the gain-rescaled couplings evaluated at the operating point, with $\boldsymbol{A}$ expressed in the source-covariance eigenbasis.

In the end, the effective noise covariance matrix that enters the subsequent mean-field calculation is therefore

$$\boldsymbol{D}_{\tilde{\boldsymbol{\eta}}} = \underbrace{\boldsymbol{D}_0}_{\text{intrinsic noise}} + \underbrace{\boldsymbol{\Lambda}^\dagger\, \boldsymbol{D}_s\, \boldsymbol{\Lambda}^{\dagger\mathsf{T}}}_{\text{structured afferent drive}}\,, \tag{24}$$

which extends the intrinsic-noise-only covariance of [1, 4] by a structured, input-driven contribution.

### 3 Source covariance geometry and moment matching

The composite noise covariance in Eq. (24) shows how afferent geometry enters the target problem. To connect the source geometry to experimentally measured quantities, we summarize the source spectrum using two observables. The first is the total source variance $\mathrm{Tr}(\boldsymbol{\Sigma}_s) = \mathrm{Tr}(\boldsymbol{D}_s) = \sum_{\alpha=1}^M \lambda_\alpha$, which measures the overall strength of the afferent fluctuations. The second is source dimensionality,

$$\Gamma_s = \frac{\mathrm{Tr}(\boldsymbol{\Sigma}_s)^2}{\mathrm{Tr}(\boldsymbol{\Sigma}_s^2)} = \frac{\mathrm{Tr}(\boldsymbol{D}_s)^2}{\mathrm{Tr}(\boldsymbol{D}_s^2)} = \frac{\left(\sum_{\alpha=1}^M \lambda_\alpha\right)^2}{\sum_{\alpha=1}^M \lambda_\alpha^2}\,, \tag{25}$$

which measures the effective dimensionality of source-area covariance. If all source eigenvalues are equal, variance is spread uniformly across modes and $\Gamma_s$ approaches its maximum value $M$. If variance is concentrated in a few large eigenvalues, $\Gamma_s$ is smaller, indicating that source activity occupies fewer effective dimensions. Therefore, $\mathrm{Tr}(\boldsymbol{\Sigma}_s)$ quantifies how much source variance is available to drive the target, whereas $\Gamma_s$ quantifies how many independent source directions carry that variance. Our goal is to express the downstream covariance calculation in terms of these two source-level observables. We therefore model the eigenvalues $\lambda_\alpha$ as independent draws from a common distribution and choose the parameters of that distribution so that its first two moments match the measured $\mathrm{Tr}(\Sigma_s)$ and $\Gamma_s$. This matching is justified because the number of afferent inputs $M$ is large enough so that we can replace the sums over source eigenvalues above by their average. This yields

$$\mathbb{E}[\lambda_\alpha] = \frac{\mathrm{Tr}(\boldsymbol{\Sigma}_s)}{M}\,, \qquad \mathbb{E}[\lambda_\alpha^2] = \frac{\mathrm{Tr}(\boldsymbol{\Sigma}_s)^2}{M\Gamma_s}\,, \tag{26}$$

for the first two moments of the eigenvalue distribution. These two identities hold generally for any distribution of $\lambda_\alpha$. For concreteness and numerical simulations, we discuss in the following two reasonable example distributions of eigenvalues.

**Lognormal source spectrum.** A natural choice for positive mode variances is a lognormal distribution

$$\lambda_\alpha \sim \mathcal{LN}(\mu_\lambda, \sigma_\lambda^2)\,, \tag{27}$$

with moments

$$\mathbb{E}[\lambda_\alpha] = e^{\mu_\lambda + \sigma_\lambda^2/2}\,, \qquad \mathbb{E}\big[\lambda_\alpha^2\big] = e^{2\mu_\lambda + 2\sigma_\lambda^2}\,. \tag{28}$$

The lognormal model allows the source spectrum to be strongly heterogeneous, with a small number of modes potentially carrying large variance. Matching the expected total source variance gives

$$\mathrm{Tr}(\boldsymbol{\Sigma}_s) = \mathrm{Tr}(\boldsymbol{D}_s) \approx M\,\mathbb{E}[\lambda_\alpha] = M\,e^{\mu_\lambda + \sigma_\lambda^2/2}\,, \tag{29}$$

while matching the expected source dimensionality gives

$$\Gamma_s \approx \frac{(M\,\mathbb{E}[\lambda_\alpha])^2}{M\,\mathbb{E}[\lambda_\alpha^2]} = M\,e^{-\sigma_\lambda^2}\,. \tag{30}$$

Together, these two constraints determine the lognormal distribution

$$\mu_\lambda = \ln\!\left(\frac{\mathrm{Tr}(\boldsymbol{\Sigma}_s)}{M}\right) - \frac{1}{2}\ln\!\left(\frac{M}{\Gamma_s}\right)\,, \qquad \sigma_\lambda^2 = \ln\!\left(\frac{M}{\Gamma_s}\right)\,. \tag{31}$$

Thus, once the total source variance and dimensionality are specified, the corresponding lognormal source spectrum is fixed.

**Gamma source spectrum.** As an alternative positive distribution, we can model the source eigenvalues using a Gamma distribution

$$\lambda_\alpha \sim \mathrm{Gamma}(k, \theta)\,, \tag{32}$$

with moments

$$\mathbb{E}[\lambda_\alpha] = k\theta\,, \qquad \mathbb{E}\big[\lambda_\alpha^2\big] = k(k+1)\theta^2\,. \tag{33}$$

The same moment-matching procedure gives $\mathrm{Tr}(\boldsymbol{\Sigma}_s) \approx M\,k\theta$ and $\Gamma_s \approx kM/(k+1)$, from which

$$k = \frac{\Gamma_s}{M - \Gamma_s}\,, \qquad \theta = \frac{M - \Gamma_s}{M\,\Gamma_s}\,\mathrm{Tr}(\boldsymbol{\Sigma}_s)\,. \tag{34}$$

The Gamma distribution provides a second parametrization of positive source spectra. Unlike the lognormal distribution, it has a different tail behavior, but this difference does not affect the leading-order calculation below because only the first two source-eigenvalue moments enter the final covariance-moment expressions.

### 4 Path-integral formulation and target dimensionality

We now compute the covariance moments required to obtain target dimensionality. The covariance $C$ in Eq. (18) depends on a particular realization of the effective recurrent connectivity $W^*$, the intrinsic noise strengths $D_{0,i}$, the source eigenvalues $\lambda_\alpha$, and the rotated projectors $\Lambda^\dagger_{i\alpha}$. We therefore seek typical covariance statistics after averaging over these sources of disorder. Throughout this section, the average over the internal disorder induced by the connectivities $W_{ij}$ is denoted by $\langle\cdot\rangle_W$ and the average over all external disorder sources by $\langle\cdot\rangle_D$. The full disorder average $\langle\cdot\rangle_{W,D}$ is abbreviated by $\langle\cdot\rangle$.

Dimensionality depends not only on the mean covariance magnitude, but also on the entry-wise dispersion of covariance values [8, 4, 7]. In particular, we require the mean diagonal covariance, the mean off-diagonal covariance, and the second moments of diagonal and off-diagonal covariance entries. These quantities are obtained using a beyond-mean-field calculation based on the moment-generating function of the zero-frequency activity fluctuations, following the formalism introduced in [1], but with the intrinsic noise covariance replaced by the composite covariance $D_{\tilde{\eta}}$ in Eq. (24). At zero frequency, the linearized dynamics imply $\tilde{\boldsymbol{x}} = (\mathbb{1} - \boldsymbol{W}^*)^{-1}\tilde{\boldsymbol{\eta}}$. The probability density of $\tilde{\boldsymbol{x}}$ can therefore be written by enforcing this linear relation with a delta function

$$p(\tilde{\boldsymbol{x}}) = \int d\tilde{\boldsymbol{\eta}}\, p(\tilde{\boldsymbol{\eta}})\, \delta\big(\tilde{\boldsymbol{x}} - (\mathbb{1} - \boldsymbol{W}^*)^{-1}\tilde{\boldsymbol{\eta}}\big) = |\det(\mathbb{1} - \boldsymbol{W}^*)| \int d\tilde{\boldsymbol{\eta}}\, p(\tilde{\boldsymbol{\eta}})\, \delta(\tilde{\boldsymbol{\eta}} - (\mathbb{1} - \boldsymbol{W}^*)\tilde{\boldsymbol{x}}) \ . \tag{35}$$

The corresponding moment-generating function is

$$\begin{aligned} Z(\boldsymbol{J}) &= \int d\tilde{\boldsymbol{x}}\, \exp\Big\{\boldsymbol{J}^\mathsf{T}\tilde{\boldsymbol{x}}\Big\}\, p(\tilde{\boldsymbol{x}}) \\ &= \frac{|\det(\mathbb{1} - \boldsymbol{W}^*)|}{(2\pi i)^N} \int d\tilde{\boldsymbol{x}}\, d\tilde{\boldsymbol{u}}\, \exp\Big\{\tilde{\boldsymbol{u}}^\mathsf{T}(\mathbb{1} - \boldsymbol{W}^*)\,\tilde{\boldsymbol{x}} + \boldsymbol{J}^\mathsf{T}\tilde{\boldsymbol{x}}\Big\} \Big\langle \exp\Big\{-\tilde{\boldsymbol{u}}^\mathsf{T}\tilde{\boldsymbol{\eta}}\Big\}\Big\rangle_{\tilde{\boldsymbol{\eta}}} \ . \end{aligned} \tag{36}$$

where the source field $\boldsymbol{J}$ generates moments of the zero-frequency target activity and, to express the delta function in exponential form, we introduced an auxiliary field $\tilde{\boldsymbol{u}}$. Averaging over the composite noise $\tilde{\boldsymbol{\eta}}$ gives

$$\Big\langle \exp\Big\{-\tilde{\boldsymbol{u}}^\mathsf{T}\tilde{\boldsymbol{\eta}}\Big\}\Big\rangle_{\tilde{\boldsymbol{\eta}}} = \exp\Big\{\frac{1}{2}\,\tilde{\boldsymbol{u}}^\mathsf{T}\,\boldsymbol{D}_{\tilde{\boldsymbol{\eta}}}\,\tilde{\boldsymbol{u}}\Big\} = \exp\Big\{\frac{1}{2}\,\tilde{\boldsymbol{u}}^\mathsf{T}\,\Big(\boldsymbol{D}_0 + \boldsymbol{\Lambda}^\dagger\,\boldsymbol{D}_s\,\boldsymbol{\Lambda}^{\dagger\mathsf{T}}\Big)\,\tilde{\boldsymbol{u}}\Big\} \ , \tag{37}$$

where we used Eq. (24). Substituting this expression into Eq. (36) gives

$$Z(\boldsymbol{J}) = \frac{|\det(\mathbb{1} - \boldsymbol{W}^*)|}{(2\pi i)^N} \int d\tilde{\boldsymbol{x}}\, d\tilde{\boldsymbol{u}}\, \exp\Big\{\tilde{\boldsymbol{u}}^\mathsf{T}(\mathbb{1} - \boldsymbol{W}^*)\,\tilde{\boldsymbol{x}} + \boldsymbol{J}^\mathsf{T}\tilde{\boldsymbol{x}} + \frac{1}{2}\,\tilde{\boldsymbol{u}}^\mathsf{T}\,\Big(\boldsymbol{D}_0 + \boldsymbol{\Lambda}^\dagger\,\boldsymbol{D}_s\,\boldsymbol{\Lambda}^{\dagger\mathsf{T}}\Big)\,\tilde{\boldsymbol{u}}\Big\} \ . \tag{38}$$

The expression above is the path-integral representation of the zero-frequency linearized network. It has the same structure as the generating function used in [1], except that the noise covariance entering the quadratic auxiliary-field term is now the composite matrix $\boldsymbol{D}_{\tilde{\boldsymbol{\eta}}} = \boldsymbol{D}_0 + \boldsymbol{\Lambda}^\dagger\,\boldsymbol{D}_s\,\boldsymbol{\Lambda}^{\dagger\mathsf{T}}$. This replacement is what inserts structured afferent geometry into the covariance-moment calculation.
The remaining steps follow the disorder-averaged covariance formalism reported in [1]. Briefly, one averages over the random recurrent matrix $\boldsymbol{W}^*$, and evaluates the resulting auxiliary-field theory around its saddle point, including the leading fluctuations required to obtain second moments of covariance entries. The result is a set of closed expressions for $\langle C_{ij}\rangle$ and $\langle C_{ij}^2\rangle$ in terms of the first two moments of the effective noise covariance $D_{\tilde{\eta}}$. Thus, in the present model, the only additional ingredients needed are $\langle D_{\tilde{\eta},ij}\rangle_D$ and $\langle D_{\tilde{\eta},ij} D_{\tilde{\eta},k\ell}\rangle_D$.
Target dimensionality is

$$D_{\mathrm{raw}}(\boldsymbol{C}) = \frac{\left(\sum_i C_{ii}\right)^2}{\sum_i C_{ii}^2 + \sum_{i\neq j} C_{ij}^2} = \frac{N\,\langle C_{ii}\rangle^2}{\langle C_{ii}^2\rangle + (N-1)\,\langle C_{ij}^2\rangle} \ . \tag{39}$$

The second equality uses self-averaging in the large-$N$ limit: sums over neurons in a typical large realization are replaced by disorder-averaged diagonal and off-diagonal covariance moments. In [1], the first and second moments of $\boldsymbol{C}$ are given by

$$\langle C_{ij}\rangle = \big[\langle \boldsymbol{D}_{\tilde{\boldsymbol{\eta}}}\rangle_D + Q^*\big(\langle \boldsymbol{D}_{\tilde{\boldsymbol{\eta}}}\rangle_D\big)\ \mathbb{1}\big]_{ij} = \big[\langle \boldsymbol{D}_{\tilde{\boldsymbol{\eta}}}\rangle_D - \mathrm{diag}\left\{(\mathbb{1} - \boldsymbol{K})\mathrm{vec}\left\{\langle \boldsymbol{D}_{\tilde{\boldsymbol{\eta}}}\rangle_D\right\}\right\}\big]_{ij} \ , \tag{40a}$$

$$\begin{aligned} \langle C_{ij}^2\rangle = (1 + \delta_{ij})\,\Big[\boldsymbol{K}\,\langle\langle \boldsymbol{C}\rangle_W \odot \langle \boldsymbol{C}\rangle_W\rangle_D\,\boldsymbol{K}^\mathsf{T}\Big]_{ij} \\ - \delta_{ij}\ \langle C_{ii}\rangle_{W,D}^2 \ , \end{aligned} \tag{40b}$$

with

$$K = \mathbb{I} + \frac{g_{\text{eff}}^2}{N(1-g_{\text{eff}}^2)}\, U_N\,, \tag{41}$$

where $U_N$ is the $N \times N$ all-ones matrix and $g_{\text{eff}}$ is the effective spectral radius introduced in Eq. (10). Operator $\operatorname{diag}\{\boldsymbol{v}\} = \boldsymbol{M}$ produces a diagonal matrix $M_{ij} = \delta_{ij} v_i$ from a vector $\boldsymbol{v}$ and operator $\operatorname{vec}\{\boldsymbol{M'}\} = \boldsymbol{v'}$ produces a vector $v'_i = M'_{ii}$ from the diagonal elements of a matrix $\boldsymbol{M'}$. The function $Q^*(\cdot)$ summarizes recurrent amplification of noise modulations at the saddle point. The covariances averaged only over the intrinsic disorder $\langle \boldsymbol{C}\rangle_W$ in Eq. (40b) can be obtained from Eq. (40a) by dropping the average over the external input $\langle\cdot\rangle_D$.

Relative to [1], $\langle D_{\tilde{\eta},ij}\rangle_D$ and $\langle D_{\tilde{\eta},ij} D_{\tilde{\eta},k\ell}\rangle_D$ are modified by structured afferent drive in Eq. (24); we therefore require the first two moments of the composite noise covariance.

We model the intrinsic noise variances $D_{0,i}$ as independent lognormal random variables,

$$D_{0,i} \sim \mathcal{LN}\left(\ln D_0 - \frac{1}{2}\ln\left(\frac{D_0^2+\Delta_0}{D_0^2}\right),\, \ln\left(\frac{D_0^2+\Delta_0}{D_0^2}\right)\right)\,. \tag{42}$$

The lognormal parametrization ensures positive intrinsic variances while allowing neuron-to-neuron heterogeneity. The parameter $D_0$ controls the mean intrinsic fluctuation strength, whereas $\Delta_0$ controls its dispersion across target neurons. We next compute the moments of the composite covariance. Averaging Eq. (24) over $\boldsymbol{\Lambda}^\dagger$ and $\boldsymbol{D}_s$, while treating source eigenvalues and projectors as independent, gives

$$\langle D_{\tilde{\eta},ij}\rangle_D = \delta_{ij}\, D_0 + \sum_{\alpha=1}^{M} \mathbb{E}[\lambda_\alpha]\, \mathbb{E}\left[\Lambda^\dagger_{i\alpha}\,\Lambda^\dagger_{j\alpha}\right]\,. \tag{43}$$

Using the Gaussian afferent-coupling model in Eq. (23), $\mathbb{E}[\Lambda^\dagger_{i\alpha}\Lambda^\dagger_{j\alpha}] = \delta_{ij} V$. Combining this with the first source-spectrum moment in (26) gives

$$\langle D_{\tilde{\eta},ij}\rangle_D = \delta_{ij}\left(D_0 + \operatorname{Tr}(\boldsymbol{\Sigma}_s)\, V\right)\,. \tag{44}$$

Thus, the mean composite noise covariance is diagonal. On average, structured afferent drive increases the variance of each target neuron by the total source variance multiplied by the afferent coupling variance $V$. This mean effect depends on $\operatorname{Tr}(\Sigma_s)$, but not on source dimensionality $\Gamma_s$.

Dimensionality also depends on the dispersion of covariance entries, so we need the second moment of $D_{\tilde{\eta}}$. Subtracting the product of means gives

$$\begin{aligned}\langle D_{\tilde{\eta},ij}\, D_{\tilde{\eta},k\ell}\rangle_D - \langle D_{\tilde{\eta},ij}\rangle_D\,, \langle D_{\tilde{\eta},k\ell}\rangle_D = {}& \delta_{ij}\,\delta_{k\ell}\left(\langle D_{0,i}\, D_{0,k}\rangle_D - \langle D_{0,i}\rangle_D\, \langle D_{0,k}\rangle_D\right) \\ &+ \sum_{\alpha,\beta} \mathbb{E}[\lambda_\alpha\,\lambda_\beta]\; \mathbb{E}\left[\Lambda^\dagger_{i\alpha}\,\Lambda^\dagger_{j\alpha}\,\Lambda^\dagger_{k\beta}\,\Lambda^\dagger_{\ell\beta}\right] \\ &- \sum_{\alpha,\beta} \mathbb{E}[\lambda_\alpha]\; \mathbb{E}[\lambda_\beta]\; \mathbb{E}\left[\Lambda^\dagger_{i\alpha}\,\Lambda^\dagger_{j\alpha}\right]\; \mathbb{E}\left[\Lambda^\dagger_{k\beta}\,\Lambda^\dagger_{\ell\beta}\right]\,. \end{aligned} \tag{45}$$

The first term captures heterogeneity in intrinsic target noise. The remaining terms capture variability introduced by the random projection of source modes into the target population. Because the rotated projectors are zero-mean Gaussian variables and independent across source modes, Wick's theorem gives

$$\mathbb{E}\left[\Lambda^\dagger_{a\alpha}\,\Lambda^\dagger_{b\alpha}\,\Lambda^\dagger_{c\beta}\,\Lambda^\dagger_{d\beta}\right] = \delta_{\alpha\beta}\,\left(\delta_{ac}\,\delta_{bd} + \delta_{ad}\,\delta_{bc}\right) V^2 + \delta_{ab}\,\delta_{cd}\, V^2\,. \tag{46}$$

We now have all the ingredients to compute (40). The first moments yield

$$\langle C_{ii}\rangle = \frac{D_0 + M\,\mathbb{E}[\lambda_\alpha]\, V}{1-g_{\text{eff}}^2}\,, \qquad \langle C_{ij}\rangle = 0\,. \tag{47}$$

The diagonal covariance grows with the total effective noise strength $D_0 + M\mathbb{E}[\lambda_\alpha]V$ and is amplified by recurrent feedback through the factor $1/(1-g_{\text{eff}}^2)$. The mean off-diagonal covariance vanishes by symmetry because the recurrent connectivity and rotated projectors have zero mean. The second-moment combination that enters the denominator of dimensionality is

$$\begin{aligned}\langle C_{ii}^2\rangle + (N-1)\,\langle C_{ij}^2\rangle &= \big(N + g_{\text{eff}}^2(2-g_{\text{eff}}^2)\big)\,(D_0 + M\,\mathbb{E}[\lambda_\alpha]\,V)^2 \\ &+ \big(N + g_{\text{eff}}^2(2-g_{\text{eff}}^2)\big)\,(1-g_{\text{eff}}^2)^2\Big[\Delta_0 + MV^2\,\Big((N+1)\,\mathbb{E}[\lambda_\alpha]^2 + (N+2)\,\mathrm{Var}[\lambda_\alpha]\Big)\Big]\,.\end{aligned} \tag{48}$$

This expression contains two conceptually distinct contributions. The first is set by the square of the mean effective noise strength. The second depends on heterogeneity in the composite noise covariance, including intrinsic-noise heterogeneity $\Delta_0$ and source-driven heterogeneity from the random projection of source modes into the target population. The latter contribution depends on both the first and second moments of the source eigenvalue distribution.
In the large-$N$ limit, we employ

$$N + g_{\text{eff}}^2(2-g_{\text{eff}}^2) \approx N\,, \tag{49}$$

together with $N+1 \approx N$ and $N+2 \approx N$, so that

$$(N+1)\,\mathbb{E}[\lambda_\alpha]^2 + (N+2)\,\mathrm{Var}[\lambda_\alpha] \approx N\,\mathbb{E}\big[\lambda_\alpha^2\big]\,. \tag{50}$$

Substituting into (48) yields

$$\langle C_{ii}^2\rangle + (N-1)\,\langle C_{ij}^2\rangle \approx N\,\Big\{(D_0 + M\,\mathbb{E}[\lambda_\alpha]\,V)^2 + (1-g_{\text{eff}}^2)^2\,\big[\Delta_0 + MV^2\,N\,\mathbb{E}\big[\lambda_\alpha^2\big]\big]\Big\}\,. \tag{51}$$

Combining (47) and (51) with (39) gives the compact closed-form normalized dimensionality reported in the main text:

$$D(\boldsymbol{C}) \equiv D_{\text{raw}}(\boldsymbol{C})/N = \frac{(1-g_{\text{eff}}^2)^2\,\left(D_0 + M\,\mathbb{E}[\lambda_\alpha]\,V\right)^2}{(D_0 + M\,\mathbb{E}[\lambda_\alpha]\,V)^2 + (1-g_{\text{eff}}^2)^2\,(\Delta_0 + N\,MV^2\,\mathbb{E}[\lambda_\alpha^2])}\,. \tag{52}$$

Using the moment-matching identities in (26), this can be rewritten directly in terms of source total variance and dimensionality:

$$D(\boldsymbol{C}) = \frac{(1-g_{\text{eff}}^2)^2\,\left(D_0 + \mathrm{Tr}(\boldsymbol{\Sigma}_s)\,V\right)^2}{(D_0 + \mathrm{Tr}(\boldsymbol{\Sigma}_s)\,V)^2 + (1-g_{\text{eff}}^2)^2\left(\Delta_0 + N\,V^2\,\frac{\mathrm{Tr}(\boldsymbol{\Sigma}_s)^2}{\Gamma_s}\right)}\,. \tag{53}$$

To recover the compact notation used in the main text, define

$$H \equiv \frac{\Delta_0}{D_0^2}, \qquad V_A \equiv V, \qquad V_s \equiv \frac{\mathrm{Tr}(\boldsymbol{\Sigma}_s)}{D_0}, \qquad D_s \equiv \Gamma_s.$$

Factoring $D_0^2$ from Eq. (53) then gives

$$\begin{aligned} D(\boldsymbol{C}) &= \mathcal{D}\,(1-g_{\text{eff}}^2)^2, \\ \mathcal{D}^{-1} &= 1 + (1-g_{\text{eff}}^2)^2\,\frac{H + NV_A^2V_s^2/D_s}{(1+V_AV_s)^2}, \end{aligned}$$

which is the form reported in the main text. The expression in Eq. (53) is our central theoretical result. It separates the effects of target recurrence, intrinsic target variability, source variance, and source dimensionality. The numerator is controlled by the squared mean covariance scale, amplified by the recurrent factor $(1-g_{\text{eff}}^2)^2$. The denominator contains the same mean scale plus heterogeneity terms that reduce dimensionality by concentrating covariance into fewer effective directions.

The role of source dimensionality is explicit. At fixed total source variance $\mathrm{Tr}(\mathbf{\Sigma}_s)$, increasing $\Gamma_s$ decreases $\mathbb{E}[\lambda_\alpha^2]$, meaning that source variance is distributed more evenly across modes. This reduces the source-driven heterogeneity term in the denominator and increases target dimensionality. Conversely, when $\Gamma_s$ is small, a few source modes dominate the afferent drive, increasing covariance anisotropy in the target and reducing $D(\boldsymbol{C})$.

The role of source variance is distinct. Increasing $\mathrm{Tr}(\mathbf{\Sigma}_s)$ increases the mean afferent drive entering the target, but it also increases the source-driven second-moment term. In the input-dominated regime, the latter effect concentrates shared variability into structured directions and can reduce target dimensionality. Thus, source variance and source dimensionality have separable effects: source variance controls the strength of afferent drive, whereas source dimensionality controls how many independent directions carry that drive.

Together, Eq. (53) formalizes the input-geometry mechanism. Target dimensionality is not determined solely by local recurrence or intrinsic noise: it also depends on how much variance is present in the source population and how that variance is distributed across source covariance modes. This provides the theoretical basis for the main-text prediction that source variance and source dimensionality can exert dissociable effects on downstream population geometry.

### 5 Limiting cases and Poisson-like variability

We first consider the limiting case in which the target population receives no structured afferent drive. Setting $V = 0$ in Eq. (53) removes the source-to-target contribution, leaving a recurrent target network driven only by intrinsic noise. Dimensionality then reduces to

$$D(\boldsymbol{C}) = \frac{(1-g_{\text{eff}}^2)^2\, D_0^2}{D_0^2 + (1-g_{\text{eff}}^2)^2\, \Delta_0}\,. \tag{54}$$

This expression recovers the intrinsic-noise limit of the recurrent covariance theory and provides a baseline against which the structured-afferent case can be compared.

##### Homogeneous noise model.

When intrinsic noise is homogeneous across neurons ($\Delta_0 = 0$), the denominator collapses to $D_0^2$ and

$$D(\boldsymbol{C})\big|_{\Delta_0=0} = (1-g_{\text{eff}}^2)^2\,. \tag{55}$$

This expression reproduces the findings in [5, 7].

##### Renewal-like nonlinear model.

In the following sections, we treat the linearized network model as originating from a nonlinear spiking network as in ref. [1]. Furthermore, we approximate single-neuron spike trains as stationary renewal processes [9]. In the large counting-window limit, the spike-count Fano factor $F_i$ is then equal to the squared coefficient of variation of the inter-spike intervals, $F_i = CV_{\text{ISI},i}^2$, and the zero-frequency auto-covariance satisfies

$$C_{ii} = F_i r_i^*, \tag{56}$$

where $r_i^*$ is the operating-point firing rate. The Poisson case is recovered as the special limit $F_i = 1$, while heterogeneous $F_i$ allow for more general renewal-like variability. For the linearized dynamics with effective connectivity $W^*$, the zero-frequency covariance is

$$\boldsymbol{C} = (\mathbb{1} - \boldsymbol{W}^*)^{-1}\, \boldsymbol{D}_0\, (\mathbb{1} - \boldsymbol{W}^*)^{-T}\ , \tag{57}$$

where $D_0 = \mathrm{diag}(D_{0,1}, \ldots, D_{0,N})$. Imposing the output constraint $C_{ii} = F_i r_i^*$ determines the intrinsic noise strengths:

$$D_{0,i} = \sum_j \left(\boldsymbol{B}^{-1}\right)_{ij} F_j \, r_j^* , \qquad B_{ij} = \left[(\mathbb{I} - \boldsymbol{W}^*)^{-1}\right]^2_{ij} . \tag{58}$$

Thus, $\boldsymbol{W}^*$ enters the noise calibration through the realization-dependent matrix $\boldsymbol{B}^{-1}$, complicating a direct disorder average. Following [1], we replace this exact realization-dependent calibration by a realization-independent approximation that reproduces the prescribed auto-covariances on average:

$$D_{0,i} = \sum_j (\mathbb{I} - \boldsymbol{S}^*)_{ij} \, F_j \, r_j^* , \qquad S^*_{ij} = \mathrm{Var}\big(W^*_{ij}\big) = \frac{g_{\text{eff}}^2}{N} . \tag{59}$$

This choice compensates recurrent amplification at the level of the diagonal covariance constraint, so that the disorder-averaged auto-covariances satisfy $\langle C_{ii} \rangle = F_i r_i^*$ to leading order. Defining

$$v_i \equiv F_i r_i^*, \qquad \bar{v} \equiv \frac{1}{N} \sum_j v_j , \tag{60}$$

the realization-independent noise strength becomes

$$D_{0,i} = v_i - g_{\text{eff}}^2 \bar{v}. \tag{61}$$

Because the disorder averaged covariances that appear in the participation-ratio depend only on the first two moments of $D_{0,i}$ (see Eq. (40)), we characterize $v_i$ by its mean and variance,

$$\mathbb{E}[v_i] = \bar{v} , \qquad \mathrm{Var}(v_i) = \Delta_v . \tag{62}$$

Then

$$\mathbb{E}[D_{0,i}] = (1 - g_{\text{eff}}^2) \, \bar{v} , \qquad \mathrm{Var}(D_{0,i}) = \left(1 - \frac{2 g_{\text{eff}}^2}{N} + \frac{g_{\text{eff}}^4}{N}\right) \Delta_v \xrightarrow{N \gg 1} \Delta_v . \tag{63}$$

Substituting the replacements $D_0 \to (1 - g_{\text{eff}}^2) \, \bar{v}$ and $\Delta_0 \to \Delta_v$ into Eq. (54) yields

$$D(\boldsymbol{C}) = \frac{(1 - g_{\text{eff}}^2)^2 \, \bar{v}^2}{(1 - g_{\text{eff}}^2)^2 \, \bar{v}^2 + (1 - g_{\text{eff}}^2)^2 \, \Delta_v} \times (1 - g_{\text{eff}}^2)^2 = \frac{\bar{v}^2}{\bar{v}^2 + \Delta_v} (1 - g_{\text{eff}}^2)^2 = \frac{\mathbb{E}[v_i]^2}{\mathbb{E}[v_i^2]} (1 - g_{\text{eff}}^2)^2 . \tag{64}$$

The factor $(1 - g_{\text{eff}}^2)^2$ cancels inside the diagonal-variance participation factor because the Poisson-like constraint fixes the diagonal covariances before recurrent amplification acts on them. Recurrence still reduces dimensionality through the off-diagonal covariance structure, which produces the final multiplicative factor $(1 - g_{\text{eff}}^2)^2$.
Assuming statistical independence between $F_i$ and $r_i^*$, the participation of $v_i = F_i r_i^*$ factorizes into contributions from firing-rate and Fano-factor heterogeneity, yielding

$$\begin{aligned} \frac{\mathbb{E}[v_i]^2}{\mathbb{E}[v_i^2]} (1 - g_{\text{eff}}^2)^2 &= \frac{\mathbb{E}[F_i]^2}{\mathbb{E}\big[F_i^2\big]} \times \frac{\mathbb{E}[r_i^*]^2}{\mathbb{E}[(r_i^*)^2]} \times (1 - g_{\text{eff}}^2)^2 \\ &= D(\boldsymbol{F}) \, D(\boldsymbol{r}^*) \, (1 - g_{\text{eff}}^2)^2 , \end{aligned} \tag{65}$$

where we defined Fano-factor participation

$$D(\boldsymbol{F}) \equiv \frac{\mathbb{E}[F_i]^2}{\mathbb{E}\big[F_i^2\big]} \tag{66}$$

and response participation, $D(\boldsymbol{r}^*) = \mathbb{E}[r_i^*]^2/\mathbb{E}[(r_i^*)^2]$ [10]. For Poisson neurons, $F_i = 1$ and Fano-factor participation is $D(\boldsymbol{F}) = 1$.
For an unspecified static nonlinearity $f(\cdot)$, dimensionality therefore takes the form

$$D(\boldsymbol{C}) = D(\boldsymbol{F})\, D(\boldsymbol{r}^*)\, \left(1 - g^2\, \mathbb{E}\big[(f_i^*)^2\big]\right)^2 , \tag{67}$$

This is the equation reported in the main text, where asterisks identify operating-point rates and gains and $g_{\text{eff}}$ distinguishes effective recurrence from the bare strength $g$. This expression extends Tian et al.'s result in [2] to Poisson-like neurons and to arbitrary input-output nonlinearities. The remaining task is to specify how the gain moment $\mathbb{E}\big[(f_i^*)^2\big]$ depends on the observed firing-rate distribution. Since the true synaptic input to each neuron is generally unobserved in experimental data, directly estimating the input–output function is challenging. We therefore focus on standard analytic nonlinearities that are biologically motivated, tractable, and relate the second moment of the gains to macroscopic population observables, allowing direct comparison between the theory and recorded neural activity.

**Threshold-power-law gains.** Consider the threshold-power-law nonlinearity

$$f(h) = k[h-\theta]_+^p, \qquad p > 0, \tag{68}$$

where $[x]_+ \equiv \max(x, 0)$. We assume that the neurons included in the lognormal approximation are on the active branch of the transfer function, i.e.

$$h_i^* > \theta. \tag{69}$$

Equivalently, we model the distribution of positive operating-point firing rates. On the active branch,

$$r_i^* = f(h_i^*) = k(h_i^* - \theta)^p. \tag{70}$$

The local gain is

$$f_i^* = \partial_i f(h_i^*) = kp(h_i^* - \theta)^{p-1}. \tag{71}$$

Using the fixed-point relation

$$h_i^* - \theta = \left(\frac{r_i^*}{k}\right)^{1/p}, \tag{72}$$

we can express the gain by

$$f_i^* = pk^{1/p}(r_i^*)^{(p-1)/p}.$$

After this substitution the threshold $\theta$ does not appear explicitly anymore. Once we condition on the active branch, the local gain can be written directly as a function of the observed operating-point firing rate. Thus, $\theta$ determines which neurons are active, but the gain–rate relation for active neurons depends only on $k$, $p$, and $r_i^*$. Finally we obtain

$$\mathbb{E}\big[(f_i^*)^2\big] = p^2 k^{2/p} \mathbb{E}\left[(r_i^*)^{\frac{2(p-1)}{p}}\right]. \tag{73}$$

Notably, for $p = 2$, we have $E\left[(r_i^*)\right] = \bar{r}^*$ and choosing $k = 1/4$ we recover the result in [2].
We now further assume that the positive operating-point rates are lognormally distributed,

$$r_i^* \sim \text{LogNormal}(\mu_r, \sigma_r^2), \qquad \log r_i^* \sim \mathcal{N}(\mu_r, \sigma_r^2). \tag{74}$$

Then

$$\bar{r}^* \equiv \mathbb{E}[r_i^*] = e^{\mu_r + \sigma_r^2/2}, \tag{75}$$

and

$$\mathbb{E}[(r_i^*)^2] = e^{2\mu_r + 2\sigma_r^2}. \tag{76}$$

Response participation is therefore

$$D(\boldsymbol{r}^*) = \frac{\mathbb{E}[r_i^*]^2}{\mathbb{E}[(r_i^*)^2]} = \frac{e^{2\mu_r + \sigma_r^2}}{e^{2\mu_r + 2\sigma_r^2}} = e^{-\sigma_r^2}. \tag{77}$$

For any real exponent $q$, the lognormal moment is

$$\mathbb{E}[(r_i^*)^q] = e^{q\mu_r + \frac{1}{2}q^2\sigma_r^2}. \tag{78}$$

Using $\bar{r}^* = e^{\mu_r + \sigma_r^2/2}$, we can rewrite this as

$$\begin{aligned}\mathbb{E}[(r_i^*)^q] &= \left(e^{\mu_r + \sigma_r^2/2}\right)^q e^{\frac{1}{2}q(q-1)\sigma_r^2} \\ &= (\bar{r}^*)^q e^{\frac{1}{2}q(q-1)\sigma_r^2} \\ &= (\bar{r}^*)^q D(\boldsymbol{r}^*)^{-\frac{1}{2}q(q-1)}.\end{aligned} \tag{79}$$

Substituting $q = 2(p-1)/p$, we have

$$q - 1 = \frac{2(p-1)}{p} - 1 = \frac{p-2}{p}, \tag{80}$$

and therefore

$$\frac{1}{2}q(q-1) = \frac{(p-1)(p-2)}{p^2}. \tag{81}$$

Thus,

$$\mathbb{E}\left[(r_i^*)^q\right] = (\bar{r}^*)^{2(p-1)/p} D(\boldsymbol{r}^*)^{-\frac{(p-1)(p-2)}{p^2}}. \tag{82}$$

Combining Eqs. (73) and (82), we obtain

$$\mathbb{E}\left[(f_i^*)^2\right] = p^2 k^{2/p} (\bar{r}^*)^{2(p-1)/p} D(\boldsymbol{r}^*)^{-\frac{(p-1)(p-2)}{p^2}}. \tag{83}$$

Accordingly, the effective spectral radius becomes

$$g_{\text{eff}}^2 = g^2 p^2 k^{2/p} (\bar{r}^*)^{2(p-1)/p} D(\boldsymbol{r}^*)^{-\frac{(p-1)(p-2)}{p^2}}. \tag{84}$$

Dimensionality then becomes

$$D(\boldsymbol{C}) = D(\boldsymbol{F}) D(\boldsymbol{r}^*) \left[1 - g^2 p^2 k^{2/p} (\bar{r}^*)^{2(p-1)/p} D(\boldsymbol{r}^*)^{-\frac{(p-1)(p-2)}{p^2}}\right]^2. \tag{85}$$

Several special cases are useful. For $p = 1$, the active-branch gain is constant:

$$\mathbb{E}\left[(f_i^*)^2\right] = k^2. \tag{86}$$

For $p = 2$,

$$\mathbb{E}\left[(f_i^*)^2\right] = 4k\bar{r}^*. \tag{87}$$

With $k = 1/4$, this gives

$$\mathbb{E}\left[(f_i^*)^2\right] = \bar{r}^*, \qquad g_{\text{eff}}^2 = g^2\bar{r}^*, \tag{88}$$

which recovers the effective recurrent factor used in the quadratic threshold-power-law case. For $p = 3$,

$$\mathbb{E}\left[(f_i^*)^2\right] = 9k^{2/3} (\bar{r}^*)^{4/3} D(\boldsymbol{r}^*)^{-2/9}. \tag{89}$$

**Exponential transfer function** As a second analytically tractable unbounded case, consider the exponential transfer function

$$f(h) = Ae^{\beta h}, \tag{90}$$

with $A > 0$ and $\beta > 0$. At the operating point,

$$r_i^* = Ae^{\beta h_i^*}. \tag{91}$$

The local gain is

$$f_i = \partial_i f(h_i^*) = \beta A e^{\beta h_i^*}. \tag{92}$$

Using the fixed-point relation, this becomes

$$f_i = \beta r_i^*. \tag{93}$$

Therefore,

$$f_i^2 = \beta^2 (r_i^*)^2. \tag{94}$$

Averaging over the operating-point rate distribution gives

$$\mathbb{E}\big[(f_i^*)^2\big] = \beta^2 \mathbb{E}[(r_i^*)^2]. \tag{95}$$

Under the lognormal assumption in Eq. (74),

$$\mathbb{E}[(r_i^*)^2] = \frac{(\bar{r}^*)^2}{D(\boldsymbol{r}^*)}. \tag{96}$$

Hence,

$$\mathbb{E}\big[(f_i^*)^2\big] = \beta^2 \frac{(\bar{r}^*)^2}{D(\boldsymbol{r}^*)}. \tag{97}$$

The effective spectral radius is therefore

$$g_{\text{eff}}^2 = g^2 \beta^2 \frac{(\bar{r}^*)^2}{D(\boldsymbol{r}^*)}. \tag{98}$$

In this case, dimensionality is

$$D(\boldsymbol{C}) = D(\boldsymbol{F}) D(\boldsymbol{r}^*) \left[1 - g^2 \beta^2 \frac{(\bar{r}^*)^2}{D(\boldsymbol{r}^*)}\right]^2. \tag{99}$$

This exponential case is useful as a robustness check. Like the threshold-power-law family, it is unbounded and is therefore compatible with an unbounded lognormal approximation to the firing-rate distribution. However, it weights high-firing-rate neurons more strongly than the quadratic threshold-power-law case, because the squared gain scales as $(r_i^*)^2$ rather than as $r_i^*$. Thus, if active/passive conclusions are stable under both the quadratic power-law gain and the exponential gain, they are less likely to depend on the specific nonlinear transfer function chosen (cf. Fig. S4).

#### Renewal with afferent input

We now impose the same renewal-like constraint in the presence of structured afferent input. With afferent input, part of the target variance is already supplied by the diagonal of the structured input covariance. Therefore, the intrinsic noise must be interpreted as a residual contribution

$$D_{0,i} = v_i - g_{\text{eff}}^2 \bar{v} - \left[\boldsymbol{\Lambda}^\dagger \, \boldsymbol{D}_s \, \boldsymbol{\Lambda}^{\dagger\mathsf{T}}\right]_{ii}. \tag{100}$$

For compactness, we define

$$E_{ij} \equiv \left[\boldsymbol{\Lambda}^{\dagger}\,\boldsymbol{D}_s\,\boldsymbol{\Lambda}^{\dagger\mathsf{T}}\right]_{ij}. \tag{101}$$

The composite input covariance is then

$$\begin{aligned} D_{\tilde{\boldsymbol{\eta}},ij} &= D_{0,i}\delta_{ij} + E_{ij} \\ &= \left(v_i - g_{\text{eff}}^2 \bar{v}\right)\delta_{ij} + (1-\delta_{ij})\left[\boldsymbol{\Lambda}^{\dagger}\,\boldsymbol{D}_s\,\boldsymbol{\Lambda}^{\dagger\mathsf{T}}\right]_{ij}. \end{aligned} \tag{102}$$

Thus, the diagonal afferent contribution is cancelled by the definition of $D_{0,i}$, while the off-diagonal afferent covariance remains. In other words, the renewal condition fixes the diagonal of the composite input covariance, but structured afferent input still contributes shared fluctuations between target neurons. Setting $V = 1/M$, the expectation value of the intrinsic noise now yields

$$\langle D_{0,i}\rangle_D = \left(1 - g_{\text{eff}}^2\right)\bar{v} - \bar{\sigma}_s. \tag{103}$$

The mean of the composite covariance, however, is independent of $\bar{\sigma}_s = MV\mathbb{E}[\lambda_\alpha]$:

$$\langle D_{\tilde{\boldsymbol{\eta}},ij}\rangle_D = \left(1 - g_{\text{eff}}^2\right)\bar{v}\,\delta_{ij}. \tag{104}$$

This cancellation is a direct consequence of imposing the renewal condition on the target autocovariances. Moreover, these equations highlight that $D_{0,i}$ is not an independent noise parameter in the renewal-matched afferent model. Instead, it is the residual intrinsic variance left after accounting for recurrent amplification and for the diagonal variance injected by structured afferent input. The construction therefore assumes that this residual variance is non-negative.
We next compute the second cumulant of the composite input covariance. Because $D_{0,i}$ depends explicitly on the diagonal afferent term $A_{ii}$, the intrinsic and afferent contributions are no longer statistically independent. However, after the renewal cancellation above, the composite covariance separates into a diagonal renewal term and an off-diagonal afferent term. Therefore,

$$\begin{aligned} &\langle D_{\tilde{\boldsymbol{\eta}},ij}\,D_{\tilde{\boldsymbol{\eta}},k\ell}\rangle_D - \langle D_{\tilde{\boldsymbol{\eta}},ij}\rangle_D\,\langle D_{\tilde{\boldsymbol{\eta}},k\ell}\rangle_D \\ &\quad= \delta_{ij}\delta_{k\ell}\left(\delta_{ik} - \frac{2g_{\text{eff}}^2}{N} + \frac{g_{\text{eff}}^4}{N}\right)\Delta_v \\ &\qquad + (1-\delta_{ij})\,(1-\delta_{k\ell})\,MV^2\mathbb{E}\left[\lambda_\alpha^2\right](\delta_{ik}\delta_{j\ell} + \delta_{i\ell}\delta_{jk}) \\ &\quad\overset{N\gg 1}{=} \delta_{ij}\delta_{k\ell}\delta_{ik}\Delta_v + (1-\delta_{ij})\,(1-\delta_{k\ell})\,MV^2\mathbb{E}\left[\lambda_\alpha^2\right](\delta_{ik}\delta_{j\ell} + \delta_{i\ell}\delta_{jk}). \end{aligned} \tag{105}$$

Here,

$$\Delta_v \equiv \mathrm{Var}(v_i). \tag{106}$$

The first term captures heterogeneity in the renewal target $v_i$, whereas the second term captures the off-diagonal covariance induced by structured afferent input. The disorder-averaged moments entering dimensionality in (39) become, in the large-$N$ limit,

$$\begin{aligned} \langle C_{ij}\rangle &= \delta_{ij}\mathbb{E}[v] = \delta_{ij}\bar{v}, \\ \langle C_{ii}^2\rangle &= \mathbb{E}[v^2] + 2(2m+m^2)\left\{\frac{1}{N}\mathbb{E}[v^2] + MV^2\mathbb{E}[(\lambda_\alpha)^2]\right\}, \\ \langle C_{i\neq j}^2\rangle &= MV^2\mathbb{E}[(\lambda_\alpha)^2] + (2m+m^2)\left\{\frac{1}{N}\mathbb{E}[v^2] + MV^2\mathbb{E}[(\lambda_\alpha)^2]\right\}, \end{aligned} \tag{107}$$

with $m = g_{\text{eff}}^2/(1-g_{\text{eff}}^2)$ which already appeared in $\boldsymbol{K}$.
Defining

$$D(\boldsymbol{\lambda}) \equiv \frac{\mathbb{E}\left[\lambda_\alpha\right]^2}{\mathbb{E}\left[\lambda_\alpha^2\right]} = \Gamma_s/M, \tag{108}$$

inserting (107) and neglecting corrections of $\mathcal{O}(1/N)$, dimensionality becomes

$$D(\boldsymbol{C}) = \frac{\left(1 - g_{\text{eff}}^2\right)^2 \bar{v}^2}{\mathbb{E}\left[v_i^2\right] + \frac{N}{M}\frac{\bar{\sigma}_s^2}{D(\boldsymbol{\lambda})}}. \tag{109}$$

Equivalently, since

$$D(\boldsymbol{v}) = \frac{\bar{v}^2}{\mathbb{E}\left[v_i^2\right]}, \tag{110}$$

we can write

$$D(\boldsymbol{C}) = \frac{D(\boldsymbol{v})\left(1 - g_{\text{eff}}^2\right)^2}{1 + \frac{N}{M}\left(\frac{\bar{\sigma}_s}{\bar{v}}\right)^2 \frac{D(\boldsymbol{v})}{D(\boldsymbol{\lambda})}}. \tag{111}$$

Finally, if $F_i$ and $r_i^*$ are independent across neurons, then

$$D(\boldsymbol{v}) = D(\boldsymbol{F})\, D(\boldsymbol{r}^*), \tag{112}$$

and therefore

$$D(\boldsymbol{C}) = \frac{D(\boldsymbol{F})\, D(\boldsymbol{r}^*)\left(1 - g_{\text{eff}}^2\right)^2}{1 + \frac{N}{M}\left(\frac{\bar{\sigma}_s}{\bar{F}\bar{r}^*}\right)^2 \frac{D(\boldsymbol{F})\, D(\boldsymbol{r}^*)}{D(\boldsymbol{\lambda})}}. \tag{113}$$

For Poisson variability, $D(\boldsymbol{F}) = 1$, the expression reduces to the corresponding result with only the firing-rate participation $D(\boldsymbol{r}^*)$. In the absence of afferent input, $\bar{\sigma}_s = 0$, the previous renewal-like local result in Eq. (67) is recovered.

[8] Luca Mazzucato, Alfredo Fontanini, and Giancarlo La Camera. Stimuli Reduce the Dimensionality of Cortical Activity. *Frontiers in Systems Neuroscience*, 10, February 2016. Publisher: Frontiers.

[9] Nicolas Brunel. Dynamics of sparsely connected networks of excitatory and inhibitory spiking neurons. *Journal of computational neuroscience*, 8(3):183–208, 2000.

[10] Alessandro Treves and Edmund T Rolls. What determines the capacity of autoassociative memories in the brain? *Network: Computation in Neural Systems*, 2(4):371–397, January 1991.
